# Immunoinformatic Approach for the identification of T Cell and B Cell Epitopes in the Surface Glycoprotein and Designing a Potent Multiepitope Vaccine Construct Against SARS-CoV-2 including the new UK variant

**DOI:** 10.1101/2021.03.15.435391

**Authors:** Gracy Fathima Selvaraj, Kiruba Ramesh, Padmapriya Padmanabhan, Vidya Gopalan, Karthikeyan Govindan, Aswathi Chandran, Sivasubramanian Srinivasan, Kaveri Krishnasamy

## Abstract

The emergence of a novel coronavirus in China in late 2019 has turned into a SARS-CoV-2 pandemic affecting several millions of people worldwide in a short span of time with high fatality. The crisis is further aggravated by the emergence and evolution of new variant SARS-CoV-2 strains in UK during December, 2020 followed by their transmission to other countries. A major concern is that prophylaxis and therapeutics are not available yet to control and prevent the virus which is spreading at an alarming rate, though several vaccine trials are in the final stage. As vaccines are developed through various strategies, their immunogenic potential may drastically vary and thus pose several challenges in offering both arms of immunity such as humoral and cell-mediated immune responses against the virus. In this study, we adopted an immunoinformatics-aided identification of B cell and T cell epitopes in the Spike protein, which is a surface glycoprotein of SARS-CoV-2, for developing a new Multiepitope vaccine construct (MEVC). MEVC has 575 amino acids and comprises adjuvants and various cytotoxic T-lymphocyte (CTL), helper T-lymphocyte (HTL), and B-cell epitopes that possess the highest affinity for the respective HLA alleles, assembled and joined by linkers. The computational data suggest that the MEVC is non-toxic, non-allergenic and thermostable with the capability to elicit both humoral and cell-mediated immune responses. The population coverage of various countries affected by COVID-19 with respect to the selected B and T cell epitopes in MEVC was also investigated. Subsequently, the biological activity of MEVC was assessed by bioinformatic tools using the interaction between the vaccine candidate and the innate immune system receptors TLR3 and TLR4. The epitopes of the construct were analyzed with that of the strains belonging to various clades including the new variant UK strain having multiple unique mutations in S protein. Due to the advantageous features, the MEVC can be tested *in vitro* for more practical validation and the study offers immense scope for developing a potential vaccine candidate against SARS-CoV-2 in view of the public health emergency associated with COVID-19 disease caused by SARS-CoV-2.

## Introduction

SARS-CoV-2 outbreak began during late December 2019 in Wuhan, Hubei, China. The disease caused by SARS-CoV-2 was designated as coronavirus disease 2019 (COVID-19) by the World Health Organization (WHO) (Benvenuto, 2020) and WHO further declared COVID-19 pandemic on 11 March, 2020. Globally, on 5^th^ January 2021, there have been 83,326,479 confirmed cases of COVID-19, including 1,831,703 deaths according to WHO weekly epidemiological report. About 220 countries around the world are affected and more than 99.99% of the global SARS-CoV-2 cases are currently outside the China. Emergence of a new variant virus in UK (VUI 202012/01) during December, 2020 which has 70% high transmission potential when compared to the circulating strains has spread to 40 therapeutics and prophylactics (WHO Weekly Epidemiological Update).

Until now, there has been no efficacious antivirals and vaccines for treatment and prevention respectively to regulate its spread in spite of the promising attempts by researchers across the world (Heymann, 2020; Huang *et al*., 2020). The increasing rate of COVID-19 disease and the associated high morbidity and mortality emphasize the need of developing specific and safe anti-SARS-COV-2 drugs and vaccine candidate. Plethora of genomes of SARS-CoV-2 originated from various countries are being analyzed, characterized and further designated with various clades based on defined sets of mutations. There is an urgent need to craft vaccines for reinforcing immune defense against the virus including the new variant UK strain having multiple unique mutations in the Spike protein.

Immunoinformatics, a branch of bioinformatics, deals with the prediction and characterization of potential T and B cell epitopes useful for vaccine design including the development of vaccine against SARS-CoV (Yin, 2016; Slathia, 2018). The spike glycoprotein of SARS-CoV-2, which protrudes from the viral envelope, plays an important role in virulence and infection due to its high binding affinity towards ACE receptor expressed by various human epithelial cells and thereby facilitates entry of the virus into cell (Yan *et al*., 2020). The cell tropism behaviour of the virus is mainly due to this structural protein forming a characteristic crown of the virus. Besides, the new variant of SARS-CoV-2 that originated at UK during December, 2020 has 9 unique mutations in the S protein with the growing concern that whether the vaccines under trials would efficiently work against all the variants of SARS-CoV-2 (BMJ2020; 371:m4857). Hence, the S protein can be considered as a major target for vaccine development (Chung M, 2020) and the protein sequences of spike glycoprotein are explored systematically using multiple immunoinformatic-based tools, to identify various epitopes for developing an effective vaccine.

Development of multiepitope vaccine constructs (MEVC) have the advantages such as safety, chemical stability, selective activation of immune responses against the specific epitope covering both arms of immunity, and non-requirement of time consuming *in vitro* culture of viruses and a complete protein (Purcell, 2007; Dudek, 2010; Testa and Philip, 2012; Patronov and Doytchinova, 2013; Zheng *et al*., 2017). Though the immunological aspects of SARS-COV-2 are largely unknown, immunoinformatic based approaches using the sequences of targets of vaccine candidate viruses not only profoundly facilitate investigations on the immunogenic epitopes of SARS-COV-2 but also support the design of efficient peptide vaccine design. Reports have suggested that both humoral and cellular immunity play important roles in protective responses against this virus (Yang *et al*., 2004; Deming *et al*., 2007). T-cell epitopes are short peptide fragments ranging from 8 to11 amino acid residues, whereas the B-cell epitopes can have 15 to 25 residues. The experimental design and production of multi-epitope vaccines against viral diseases have improved dramatically in recent years with the advances in peptide vaccine research and immunoinformatic approaches. The design of a multi-epitope vaccine depends on the identification and assembly of B and T-cell epitopes that are capable of stimulating the immune system and thereby inducing more potent and effective both arms of immune responses.

In this current study, we predicted the CTL, HTL and B cell epitopes of spike protein from an isolate of our study and analysed the conservancy and other immunological properties with respect to various Indian and global strains representing all clades including the new variant SARS-CoV-2 isolate emerged in UK during December, 2020 through immunoinformatic tools. The analyses was performed using Spike proteins after downloading the sequences of whole genome sequences of SARS-CoV-2 isolates deposited in GISAID. We also investigated the population coverage of B and T cell epitopes from various countries affected by COVID-19. The interactions between the epitopes and their corresponding alleles were studied. Subsequently, the multi-epitope vaccine construct was designed and its biological activity was assessed by bioinformatic tools using the interaction between the vaccine candidate and the innate immune system receptors TLR3 and TLR4. We strongly believe that the outcome of the present report will support the development of a potential vaccine candidate against all SARS-CoV-2 variants.

## 2. Materials and Methods

### 2.1. Sequencing and sequences retrieval

Clinical samples suspected of SARS-CoV-2 infection were tested in King Institute of Preventive Medicine and Research, Chennai, Tamilnadu, India by Real Time RT PCR using TaqPath Multiplex Combo kit (Thermofisher) that detects three targets such as Spike (S), Nucleocapsid (N) and ORF 1ab sequence of SARS-CoV-2. The RNA of SARS-CoV-2 positive samples were purified and sequenced at CSIR-Centre for Cellular and Molecular Biology, Secunderabad, Telangana, India using the methods described earlier (Banu *et al*., 2020) and the complete genome sequences were submitted to GISAID (https://www.gisaid.org/) database (Elbe, 2017). For the current study, 31 sequences of full-length genome sequences of SARS-CoV-2 including 2 sequences of UK variant strains were downloaded from GISAID database and the nucleotide sequences of spike protein were selected and translated by the tool Translate (https://www.expasy.org/) and were subjected to multiple sequence alignment by MEGA 10 (www.megasoftware.net). The translated amino acid sequences of spike protein were compared with the Wuhan, China (Wuhan hu-1) reference strain sequence (NC_045512.2) as well as GISAID reference strain (EPI_ISL_402124) and the mutations specific to the SARS-CoV-2 viral isolates were identified. The Maximum Likelihood method was employed to draw phylogenetic tree.

### 2.2. Selection of protein for epitope prediction

The amino acid sequence of spike protein of HCoV-19/India/CCMB_C17/2020 (GISAID accession ID: EPI_ISL_458044) was selected from the above-mentioned sequences of SARS-CoV-2 isolates for the prediction of T cell and B cell epitopes and for multi epitope vaccine construct design.

### 2.3. Prediction of physiochemical properties, antigenicity and allergenicity

The Expasy ProtParam tool was used to evaluate the physical and chemical properties of target proteins (Gasteiger, 2003). Allergenicity and antigenicity of proteins were checked through AllerTOP v2.0 (https://www.ddg-pharmfac.net/AllerTOP/method.html) (Dimitrov, 2013) and VaxiJen v2.0 (http://www.ddg-pharmfac.net/vaxijen/VaxiJen/VaxiJen.html) (Doytchinova, 2007) respectively. For antigenicity prediction of the proteins of SARS-CoV-2 with higher accuracy, Virus model available at the VaxiJen server with a threshold of 0.4 was utilized.

### 2.4. T cell Epitope Prediction

#### 2.4.1. Cytotoxic T cell epitope prediction

The NetCTL 1.2 (http://www.cbs.dtu.dk/services/NetCTL/) server was used for the prediction of cytotoxic T-lymphocyte epitopes from the protein sequence. NetCTL uses NetMHC server’s artificial neural networks (ANNs) to predict MHC binding, NetChop-3.0 to predict C-terminal cleavages and a weight matrix to calculate TAP transport efficiency to generate predictions (Larsen 2005, 2007). The NetCTL 1.2 server can predict CD8+ T-cell epitopes for 12 human leucocyte antigens (HLA supertypes such as A1, A2, A3, A24, A26, B7, B8, B27, B39, B44, B58, and B62) taking into account the proteasomal C terminal cleavage, MHC class I binding, and TAP transport efficiency. These supertypes account for the majority of human leukocyte antigen A (HLA A) (76.5%) and HLA B (67.1%) distribution (Rodrigo, 2015). In this study, the threshold value for epitope identification was set at 1.25 which has a sensitivity and specificity of 0.54 and 0.993, respectively (Utpal Kumar Adhikari, 2018). The weight on C terminal cleavage and TAP transport efficiency were used as default parameters.

#### 2.4.2. Antigenicity, Allergenicity and Toxicity

Antigenicity of the predicted epitopes were calculated by VaxiJen v2.0 (http://www.ddg-pharmfac.net/vaxijen/VaxiJen/VaxiJen.html) with a threshold of 0.4, Allergenicity was calculated by AllerTop V2.0 (https://www.ddg-pharmfac.net/AllerTOP/method.html) and the Toxicity was predicted by ToxinPred (http://crdd.osdd.net/raghava/toxinpred/) (Gupta, 2015). Only the epitopes with VaxiJen score ≥ 0.4 were predicted to be non-toxic and non-allergens, and were taken for further screening.

#### 2.4.3. Epitope Conservancy and Immunogenicity Prediction

For epitope conservancy and immunogenicity prediction, the conservancy (http://tools.iedb.org/conservancy/) (Bui, 2007) and immunogenicity (http://tools.iedb.org/immunogenicity/) (Calis, 2013) prediction tools of IEDB were used. The conservancy levels were obtained by searching for identities in the given protein sequences. This tool calculates the degree of conservancy of an epitope within a given protein sequence set at different degrees of sequence identity. The epitope with high immunogenicity was selected for further analysis.

#### 2.4.4. Allele selection

Based on the antigenicity, allergenicity, toxicity, conservancy and immunogenicity, epitopes were shortlisted from the predicted epitopes by NetCTL. For the identification of both frequently and less frequently occurring MHC-I-binding alleles, the epitopes were analyzed by the stabilized matrix base method (SMM) in the IEDB analysis tool (http://tools.iedb.org/mhci/) (Peters, 2005). The amino acid length of peptide (9 residues) and the IC_50_ value less than 100 were selected as parameters for the identification of MHC-I-binding alleles. Here, peptides with IC_50_ values < 50 nM are considered as high affinity, < 500 nM intermediate affinity, and < 5000 nM low affinity (Adhikari, 2018).

### 2.5. Helper T cell epitope prediction

For the prediction of helper T lymphocyte (HTL) epitopes, the IEDB MHC II server (http://tools.iedb.org/mhcii/) was used (Zhang, 2008). Highest immunogenic epitopes of 15-mer were selected based on their percentile rank and IC_50_ values. The SMM-align method was used for the prediction of MHC-II-binding alleles of the best candidate epitope (Nielsen, 2007). The server provides the epitope predictions for three human MHC class II isotypes that includes HLA-DR, HLA-DP and HLA-DQ. The percentile rank is given after comparing the peptides score with million 15-mers from the SWISSPROT database. The lower percentile value means a higher binding affinity of MHC-II. Only the epitopes with rank ≤1 were taken for further analysis.

#### 2.5.1. Antigenicity, Allergenicity and Toxicity

The predicted HTL epitopes were subjected to further analysis of Antigenicity by VaxiJen v2.0 (http://www.ddg-pharmfac.net/vaxijen/VaxiJen/VaxiJen.html), Allergenicity by AllerTop V2.0 (https://www.ddg-pharmfac.net/AllerTOP/method.html) and the Toxicity was predicted by ToxinPred (http://crdd.osdd.net/raghava/toxinpred/). The epitopes predicted with good antigenic values more than threshold 0.4, non-toxic and non-allergen were selected for further study.

### 2.6. Population coverage

The expression and distribution of HLA alleles vary depending on the world’s ethnicities and regions, thereby impacting the effective production of an epitope-based vaccine (Adhikari, 2018). The population coverage analysis helps in finding the efficacy of epitopes on HLA allele distribution across different populations in the world. Every epitope and its binding HLA alleles were added, and different geographic areas were also selected for this analysis. The population coverage was calculated using the IEDB population coverage tool (http://tools.iedb.org/population/), and for this purpose, selected MHC class I and MHC class II epitopes and corresponding HLA-binding alleles were considered. This tool estimates population coverage for each epitope for various regions of the world based on the distribution of human alleles that bind to MHC (Bui, 2006).

### 2.7. Three-Dimensional Structure Design of the Epitope

The shortlisted CTL and HTL epitopes were submitted for modeling to the PEP-FOLD3 peptide structure prediction server (http:// bioserv.rpbs.univ-paris-diderot.fr/services/PEP-FOLD3/) (Thévenet, 2012; Shen, 2014; Lamiable, 2016). This is a *de novo* approach aimed at predicting peptide structures from amino acid sequences. The input sequence file was submitted in FASTA format. We used the best model for analysis of the interaction with MHC-I binding alleles.

### 2.8. Molecular interaction of the epitopes with MHC I and MHC II

In order to estimate the binding affinities between the epitopes and the molecular structure of MHC I and MHC II, *in silico* molecular docking was used. The predicted structure of CTL and HTL epitopes were subjected to dock with the MHC I and II molecules. The MHC I (PDB ID: 4NT6) and MHC II (PDB ID : 4MDI) structures were downloaded from RCSB PDB database (https://www.rcsb.org/) (Berman, 2000). The epitopes were docked corresponding to MHC I and II allele protein structures by PatchDock (http://bioinfo3d.cs.tau.ac.il/PatchDock/) (Schneidman-Duhovny, 2005) server. The parameters such as clustering RMSD value and the complex type were kept as 0.4 and default. Further, for refinement of the rigid body molecular docking solutions, FireDock (http://bioinfo3d.cs.tau.ac.il/FireDock/) (Mashiach, 2008) was used. The tool will give the best 10 docked confirmation based on global energy and Vander walls interactions.

### 2.9. Linear/Continuous B cell epitope prediction

B cell epitopes were predicted using the ABCpred server (http://crdd.osdd.net/raghava/abcpred/) (Saha and Raghava, 2006). This server predicts B cell epitopes using recurrent neural network algorithm. For higher accuracy, the threshold for the prediction was kept 0.90. The predicted epitopes were checked for their antigenicity using VaxiJen server, allergenicity by AllerTop, conservancy by IEDB tool and toxicity by ToxinPred server.

### 2.10. Structure prediction, refinement and validation

To identify discontinuous/ conformational B cell epitopes, the structure has to be predicted for the selected sequence. The secondary structure was predicted by SOPMA (https://npsa-prabi.ibcp.fr/cgi-bin/secpred_sopma.pl) (Geourjon,1995). The selected sequence 3D structure was predicted by SWISS-MODEL (https://swissmodel.expasy.org/) (Waterhouse, 2018) and the predicted structures were refined by 3D refine (http://sysbio.rnet.missouri.edu/3Drefine/) (Bhattacharya, 2016). The modeled structure was validated by the Ramachandran plot generated in Procheck (https://servicesn.mbi.ucla.edu/PROCHECK/) (Laskowski, 2001) and ProSA (https://prosa.services.came.sbg.ac.at/prosa.php) (Wiederstein, 2007).

### 2.11. Conformational /discontinuous B cell epitope prediction

Conformational epitopes are the most important as well as the most prevalent epitopes that are recognized by antibodies. More than 90% B cell epitopes are discontinuous as they are present in small segments on linear protein and they are brought to proximity while protein folding. All conformational epitope prediction methods require the 3D structures of proteins. Discontinuous epitopes of recombinant protein can be predicted using the ElliPro (http://tools.iedb.org/ellipro/) server at IEDB. In this tool, three algorithms implemented to determine the discontinuous B cell epitopes. 3D structure of input protein is approximated as number of ellipsoid shapes, calculate protrusion index (PI) and clusters neighboring residues. Ellipro defines PI score of each residue based on the center of mass of residue residing outside the largest possible ellipsoid. The prediction parameters like Minimum score and Maximum distance (Angstrom) were set as 0.8 and 6, respectively. ElliPro server predicts conformational and linear B cell epitopes using Thornton’s method and by MODELLER program or BLAST search of PDB predict and visualize antibody epitopes (Ponomarenko *et al*., 2007, 2008).

### 2.12. Mutation analyses of CTL, HTL and B cell epitopes across various SARS-CoV-2 isolates

To analyze the mutations across various SARS-CoV-2 isolates, about 3156 sequences of spike protein from India as well as other countries representing all clades were downloaded from GISAID and subjected to multiple sequence alignment using BioEdit 7.2 and analyzed for amino acid mutations with respect to the 36 selected CTL, HTL and B cell epitopes. Accession names of the samples and associated sequence data are given in FASTA file.

### 2.13. Designing of Multi epitope vaccine construct

For constructing a multi-epitope vaccine construct, the selected HTL, CTL and B-cell epitopes were joined by using GPGPG, AAY, EAAAK and KK linkers respectively. Five adjuvants namely, β-defensin, universal memory T cell helper peptide (TpD), PADRE (Pan HLA-DR reactive epitope), sequence, RS09TLR4 agonist and a M cell ligand were also added by using linkers into the vaccine construct. To enhance the immunogenicity, β-defensin was added to the N terminal whereas in the C terminal, M cell ligand was added which was followed by addition of HHHHHH for facilitating purification of the vaccine. For the construction of a multi-epitope vaccine against SARS-CoV-2, the method of Chauhan *et al*. (2019) was adopted with the following criteria: they (a) should be promiscuous, (b) should have overlapping CTL and HTL epitopes, (c) immunogenicity, (d) population coverage, (e) high affinity towards HLA alleles and (f) should not overlap with any human gene. Based on these criteria, the HTL and CTL epitopes were included in the final construct of the multi-epitope vaccine.

### 2.14. Antigenicity, allergenicity and physiochemical properties prediction

The antigenicity of the vaccine was determined using the VaxiJen server (http://www.ddg-pharmfac.net/vaxijen/VaxiJen/VaxiJen.html). The allergenicity of the vaccine was examined using AllerTOP v2.0 (http://www.ddg-pharmfac.net/AllerTOP/). The physiochemical characteristics of the vaccine were determined using the ProtParam tool of the ExPASy database server (http://web.expasy.org/protparam/).

### 2.15. Structure prediction, validation and docking with the receptor

The secondary structure of the subunit vaccine construct was predicted using PSIPred 4.0 Protein Sequence Analysis Workbench (http:// bioinf.cs.ucl.ac.uk/psipred/), while the tertiary structure was predicted by GalaxyWeb server (http://galaxy.seoklab.org/tbm), which is based on the TBM method (Shin *et al*., 2014). This server detects similar proteins, promote alignment with the target sequence, followed by making the model and performing model refinement. The best-modeled structure was refined using the Galaxy Refine server at (http://galaxy.seoklab.org/cgi-bin/submit.cgi?-type=REFINE) (Heo *et al*., 2013) by molecular dynamics simulation. The model of the vaccine construct with the best TMscore was validated by PROCHECK v.3.5 (https://servicesn.mbi.ucla.edu/PROCHECK/) and ProSA (https://prosa.services.came.sbg.ac.at/prosa.php) web servers. Vaccine-receptor docking was performed by the Cluspro v.2 (https://cluspro.bu.edu/) protein-protein docking web server (Kozakov D,2017) to determine the binding affinity of the vaccine with the TLR3 receptor (PDB ID:2A0Z) and TLR4 receptor (PDB ID: 3FXI). The server provides cluster scores based on rigid docking by sampling billions of conformation, pairwise RMSD energy minimization. Based on the lowest energy weight score and members, the final vaccine-Receptor complex model was selected.

### 2.16. Immune simulations of vaccine construct

To characterize the real-life immunogenic profiles and immune response of the multi-epitope vaccine, C-ImmSim server (Rapin, 2010) was utilized. Position-specific scoring matrix (PSSM) and machine learning are the two methods, based on which C-ImmSim predicts immune epitopes and immune interactions. It simultaneously simulates three different anatomical regions of mammals, *i.e.* bone marrow, thymus and tertiary lymphatic organs. The parameters like Random seed, simulation volume, and simulation step and the host HLA selection parameter for MHC class I was set on A1010, A1101, and B0702 and for DR MHC class II was set on DBR1_0101 were kept as default values and based on literature the time step to injection parameter was set to administer three injections at time steps 1, 84, and 168, corresponding to time = 0, 4 weeks, and 8 weeks with a total of 1050 simulation steps.

## 3. Results

### 3.1. Multiple sequence alignment and phylogenetic analysis

Twenty-one complete genome sequences of SARS-CoV-2 were sequenced from isolated and purified RNA after testing clinical samples as virus positive at King Institute of Preventive Medicine and Research, Chennai. These complete nucleic acid sequences were deposited and downloaded from GISAID database for the study. The multiple sequence alignment of the translated sequences of spike protein, along with spike protein of 10 other SARS-CoV-2 variant strains representing various clades, extracted from the genome sequences and analyzed with the Wuhan, China (Wuhan hu-1) reference strain (NC_045512.2) as well as GISAID reference strain (EPI_ISL_402124). A S protein sequence of hCoV-19/India/CCMB_C17/2020|EPI_ISL_458035 was selected for designing of epitopes for MEVC. The Maximum Likelihood method was employed to draw phylogenetic tree (**Fig. S1**).

### 3.2. Sequence retrieval and Prediction of physiochemical properties

For the prediction of T cell and B cell epitopes to design multiepitope vaccine construct, the amino acid sequence of spike protein HCoV-19/India/CCMB_C17/2020 was selected. The selected protein sequence was subjected to prediction of physiochemical parameters (**Table S1**). The protein has 1273 number of amino acids with the molecular weight of 141178.47 dalton. The total number of positively charged residues (Arg+Lys) and negatively charged residues (Asp+Glu) were 103 and 110 respectively. The theoretical isoelectric point (PI) was calculated as 6.24. Instability index (II) is one of the key parameters to distinguish stable and unstable protein candidates. A protein whose II < 40 is predicted as stable, and a value above 40 predicts that the protein may be unstable. The II of spike protein of SARS-CoV-2 was estimated to be 33.01, suggesting the nature of the protein. The aliphatic index of the protein was calculated as 84.67 indicating the stability of protein in wide ranging temperatures. Its Grand average of hydropathicity (GRAVY) was found to be - 0.079.

**Table 1.**
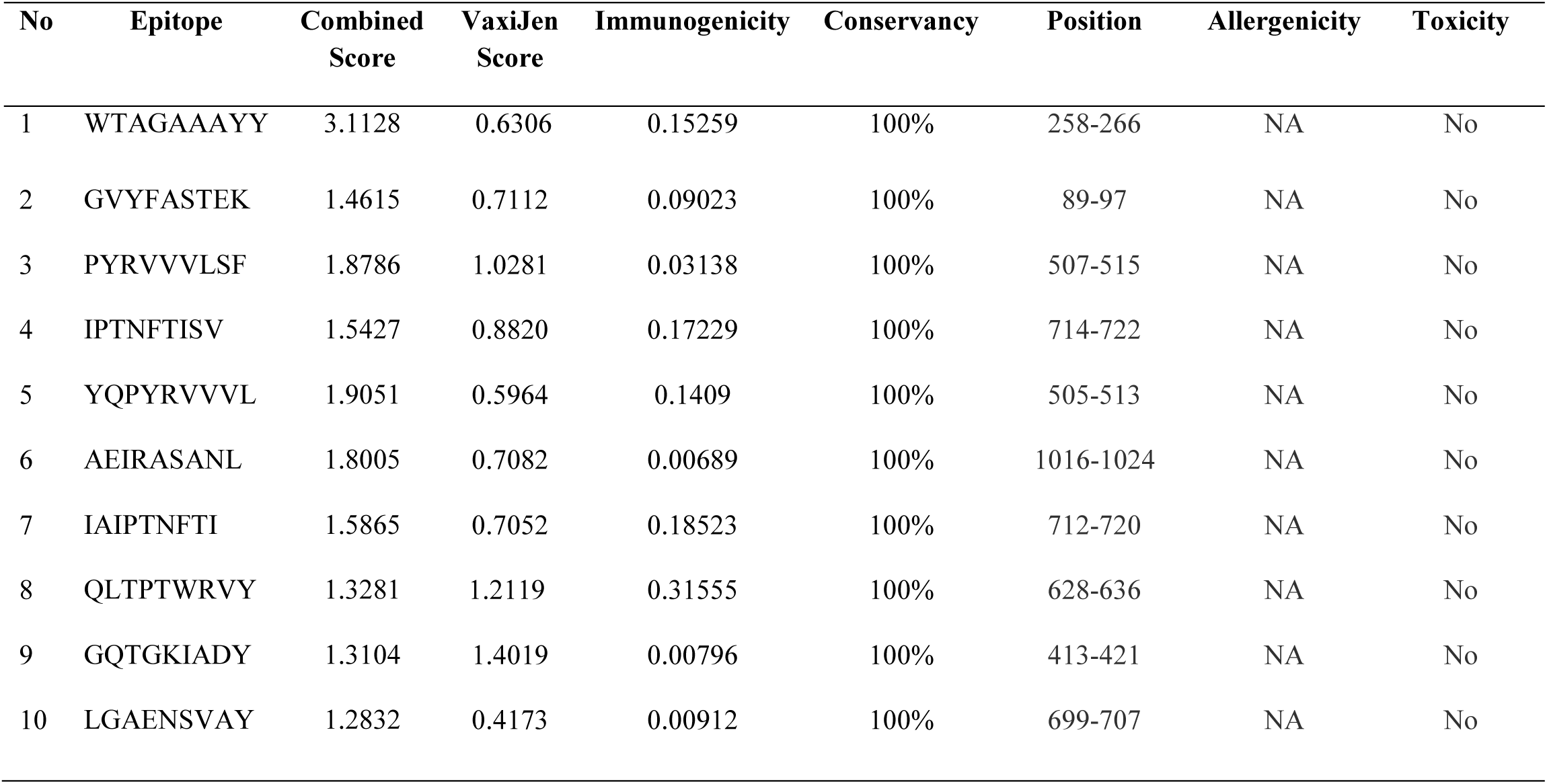
CTL epitopes with predicted features of combined score, Antigenicity, Immunogenicity, conservancy, Allergenicity and Toxicity.

### 3.3. Antigenicity and allergenicity

Prediction of the antigenicity represents a numerical criterion for the capability of the protein to bind to the B- and T-cell receptors and elicitation of immune response in the host cell. VaxiJen v2.0 was used to predict the antigenicity of the selected protein and it was observed that the protein was predicted to be potently antigenic with a value of 0.4646 at 0.4% threshold without adjuvant. Allergen proteins induce an IgE antibody response (Dimitrov *et al*., 2013) and the designed vaccine candidate must not show an allergic reaction to the body. The allergenicity of the sequence was predicted using the AllerTop tool, and the protein was predicted as non-allergen to human (**Fig. S2**).

### 3.4. T cell prediction

T-cell epitope prediction comprises identification of MHC I and MHC II binding epitopes, in order to activate both cytotoxic T-lymphocytes (CTL) and helper T-lymphocytes (HTL) mediated immune response.

#### 3.4.1. Cytotoxic T cell epitope prediction

CTL epitopes induce MHC-I cellular immune response by neutralizing virus-infected cells through releasing cytotoxic proteins. The CTL epitopes were predicted for all the selected proteins using the NetCTL 1.2 server and the combined score was considered for the epitope selection. In this study, we explored 100 epitopes in total, for which the selected threshold value was set at 1.25 (**Table S2**). The predicted T-cell epitopes were further evaluated by the VaxiJen server. It was found that 41 epitopes among the 100 primarily selected T-cell epitopes were above the threshold value 0.4 (**Table S3**), and these were subjected to immunogenicity analysis, which revealed that 21 epitopes had the immunogenicity value >0.00 (**Table S4**). From the shortlisted 21 epitopes, 10 of them showed 100% conservancy, non-toxic and non-allergen properties. The results revealed that all epitopes with the highest combined score were not the best epitopes. Among the 100 epitopes, we selected 10 epitopes, which have positive antigenicity, immunogenicity scores with 100% conservancy, non-toxic and non-allergen features (**Table 1**). These highly antigenic epitopes are predicted to interact with the MHC alleles with high affinity for eliciting an effective immune response.

**Table 2.**
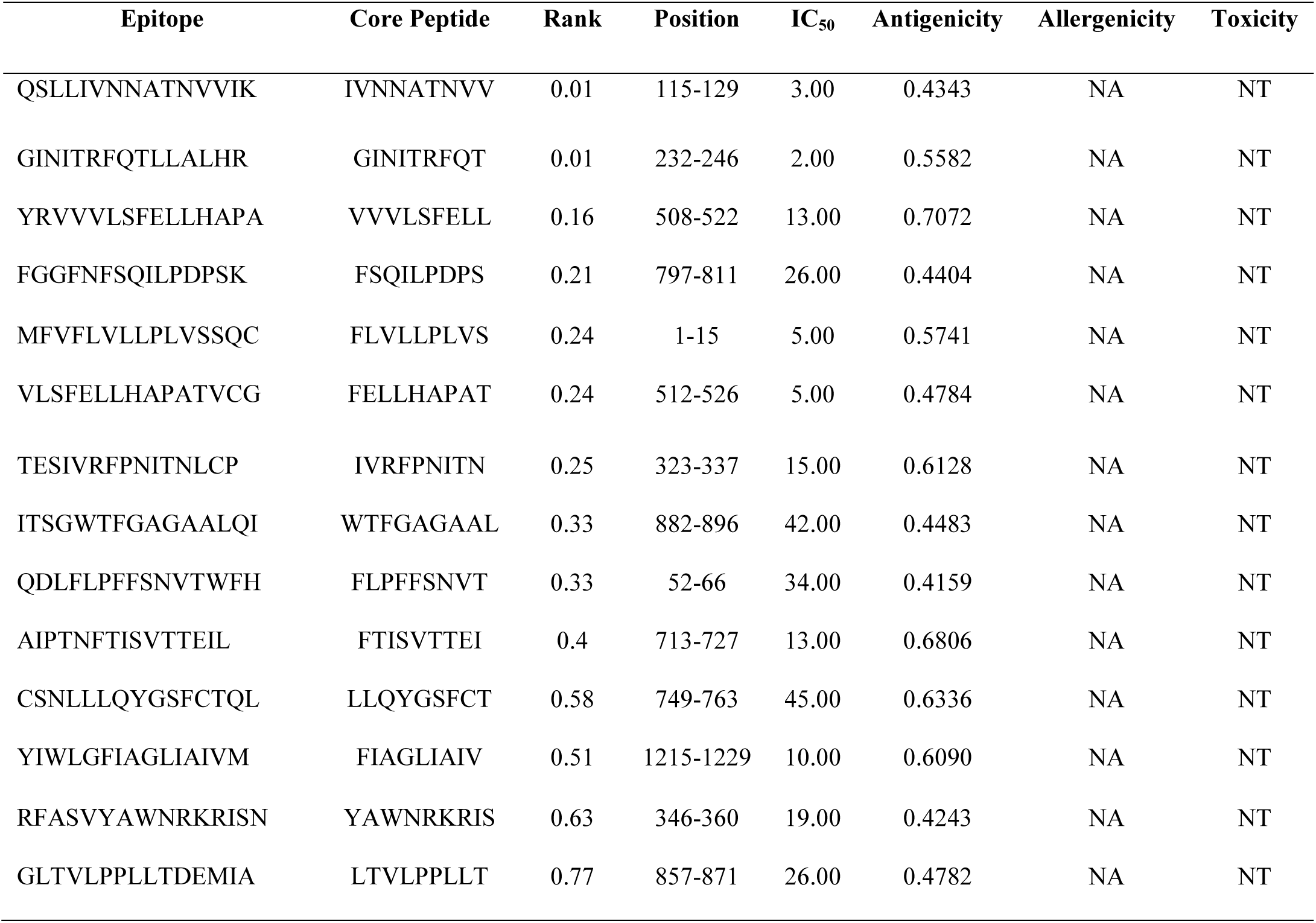
Features of Helper T cell epitopes predicted from Spike protein of SARS-CoV-2.

**Table 3.** Features of Linear or continuous B cell epitopes predicted from Spike protein of SARS-CoV-2.

| Epitope | Score | Start position | Antigenicity | Allergenicity | Toxicity | Conservancy |
| --- | --- | --- | --- | --- | --- | --- |
| GVSVITPGTNTSNQVA | 0.95 | 594 | 0.4651 | NA | NT | 100% |
| GWTAGAAAYYVGYLQP | 0.95 | 257 | 0.6210 | NA | NT | 100% |
| HRSYLTPGDSSSGWTA | 0.92 | 245 | 0.6017 | NA | NT | 100% |
| TVEKGIYQTSNFRVQP | 0.91 | 307 | 0.6733 | A | NT | 100% |
| GCLIGAEHVNNSYECD | 0.90 | 648 | 0.8480 | NA | T | 97% |
| LQSYGFQPTNGVGYQP | 0.90 | 492 | 0.5258 | NA | NT | 100% |

**Table 4.** Physiochemical properties of the MEVC.

| Features | Value |
| --- | --- |
| Number of aminoacids | 575 |
| Molecular Weight | 61480.71 |
| Theoretical pI | 9.92 |
| Total number of Negatively charged residues(Asp +Glu) | 21 |
| Total number of Positively charged residues(Arg +Lys) | 67 |
| Total number of atoms | 8752 |
| Instability Index | 26.87 |
| Aliphatic Index | 84.07 |
| Grand Average of Hydropathicity (GRAVY) | 0.001 |

Epitope conservancy among the different strains of the selected protein can provide effective immunization, and hence, the conservancy score is one of the important criteria for developing an effective vaccine. Antigenicity score plays an important role in the selection of epitopes. In this study, out of the 10 selected epitopes, 3 showed VaxiJen score > 1, one showed 0.8820, and 3 epitopes showed the values > 0.7 and the results suggested that all epitopes were highly antigenic in nature.

The immunogenicity score of the epitope can be a good criterion for the selection of the best epitope. The higher score designates a greater probability of eliciting an effective immune response. Herein, the epitope QLTPTWRVY (0.31555) is observed to be more immunogenic followed by IAIPTNFTI (0.18523), IPTNFTISV (0.17229), WTAGAAAYY (0.15259), and YQPYRVVVL (0.1409). Other epitopes such as PYRVVVLSF (0.03138), GVYFASTEK (0.09023), AEIRASANL (0.00689), GQTGKIADY (0.00796) and LGAENSVAY (0.00912) were found to be less immunogenic (**Table 1**).

#### 3.4.2. Allele selection

The selected 10 T-cell epitopes were found to be recognized by the MHC class-I molecules such as HLA-A, HLA-B, and HLA-C according to IEDB MHC class 1-binding analysis resource. We selected humans as MHC source species and the SMM method for the prediction of a distinct set of MHC HLA alleles for the humans. This tool gives an output result for HLA-binding affinity of the epitopes in the IC_50_ nM unit. A lower IC_50_ value indicates higher binding affinity of the epitopes with the MHC class I molecule. So, in this study, we chose IC_50_ values < 100 nM (IC_50_ < 100) for ensuring high affinity **(Table S5)**.

These epitopes showed affinity to varying number of HLA alleles with lower IC_50_ values. WTAGAAAYY showed the highest affinity to 7 HLA alleles followed by GVYFASTEK and AEIRASANL (5 HLA alleles); PYRVVVLSF, YQPYRVVVL and QLTPTWRVY (4 HLA alleles); IAIPTNFTI and LGAENSVAY (3 HLA alleles); GQTGKIADY (2 HLA alleles); and IPTNFTISV (1 HLA allele). Besides, all these epitopes showed the highest affinity with the allele HLA-C*12:03 with various IC_50_ values.

#### 3.4.3. Helper T cell epitope prediction

HTLs, being a key player of adaptive immune response, coordinate with B lymphocytes, TC cells and macrophages via signaling through expression of cytokines upon recognizing the epitopes presented on MHC II binding cleft. Fourteen epitopes, predicted using IEDB MHC II server, were selected based on the percentile rank (≤ 1) and less IC_50_ value **(Table 2)** indicating their higher binding affinity. Besides, these epitopes were also found to possess high antigenicity, non-allergen and non-toxic properties and were used for multi-epitope vaccine construct. They were further selected and subjected for their corresponding allele selection based on their affinity **(Table S6)**.

### 3.5. 3D structure design of the epitope and interaction of epitope with MHC I and II

MHC class I and class II alleles have a specific binding cleft where the peptide binds and are presented on the cell surface for immune system activation. Both HTL and CTL epitope structures were modelled using Pepfold-3. To examine the molecular interaction of our selected epitope with their respective HLA alleles, we performed protein-protein docking using PatchDock, for which HLA alleles and epitopes were taken as receptors and ligands respectively. The resultant docked complexes are then refined by the rigid body molecular docking solutions, FireDock **(Fig.S3 and S4).**

### 3.6. Linear/Continuous B cell epitope prediction

The prediction of B-cell epitopes from the antigenic spike protein is the major step of epitope-based vaccine design. *In silico* prediction was performed through the web server ABC pred with cutoff value of 0.90 for obtaining essential B-cell epitope candidates in the spike protein. Totally, 6 epitopes were predicted to be antigenic, and among them, TVEKGIYQTSNFRVQP epitope showed allergy to human whereas the epitope GCLIGAEHVNNSYECD displayed toxicity with 97% conservancy **(Table 3)**. The results further revealed that the four other epitopes such as GVSVITPGTNTSNQVA, GWTAGAAAYYVGYLQP, HRSYLTPGDSSSGWTA and LQSYGFQPTNGVGYQP could be the most potential B-cell epitope candidates for peptide-based vaccine design as these epitopes are conserved, antigenic, nontoxic, and non-allergenic to the human.

### 3.7. Secondary and tertiary structure prediction, refinement and validation

Identification of discontinuous/conformational B cell epitopes was based on the 3D structure of the selected spike protein sequence. The 2D structures of spike protein predicted by SOPMA tool **(Fig. S5)** showed the secondary structures such as α-helix (∼28.59%), β-turn (∼3.38%), random coil (∼44.78%) and β-pleated sheet (∼23.25%). 3D structure of the selected spike protein is not available in the PDB database, and hence 3D structure was predicted by the SWISS-MODEL server and refined by 3D Refine (**Fig. S6a**). The 3D refine score as well as GDT-TS and GDT-HA were found to be 53416.2, 1.000, and 0.9908 respectively for the selected best model. Besides, PROCHECK was used to check the stereochemical quality of the structure by considering residue geometry and overall structural geometry. The results showed 90.4% residues distributed in most favored region, 8.3% in the additional allowed region, 0.8% in the generously allowed region, and 0.5% in the disallowed region in the Ramachandran plot statistics (**Fig. S6b**). To check the potential errors of the protein 3D model, ProSA was used, and it predicted the negative Z-score of −12.71, which ensured the good quality of the model (**Fig. S6c**).

### 3.8. Conformational /discontinuous B cell epitope prediction

In this study, a total of 6 discontinuous or conformational B-cell epitopes were predicted using the ElliPro tool of IEDB. The epitopes having a protrusion index (PI) score above 0.8 were taken into consideration and the PI value of the 6 predicted epitopes ranged from 0.806 to 0.97. The residues with higher scores indicate better solvent accessibility. The predicted 3D structure was submitted for conformational epitope prediction. Conformational epitopes and their individual residues, residue position, length, and the scores are shown in **Table S7**, whereas the positions of epitopes on 3D structures are displayed in **Figs. S7 (a-f).**

### 3.9. Mutation analyses of epitopes across various SARS-CoV-2 isolates

Totally 3156 sequences of spike protein from India as well as other countries representing all clades (clade O: 686; clade S: 81; clade L: 23; clade V: 3; clade G: 662; clade GH: 564; and clade GR: 1,137) including 2 UK variant strains that fall under the clade GR were downloaded from GISAID (**Supplementary File A**), aligned and analyzed for amino acid mutations with respect to the 36 selected CTL, HTL and B cell epitopes (**Supplementary File B**). Our results indicated that all the CTL, HTL and B cell peptide epitopes were highly conserved across the all 3156 spike proteins that were subjected to analyses and the epitopes showed homology in the range of 97.53% to 100%. CTL, HTL and B cell (Linear) and B cell (discontinuous) epitopes showed homology in the range of 99.4% to 100%, 97.85% to 100% and 97.53% to 100% and 99.84 to 100% respectively. There are only 3 distinct substitution mutations identified in one of the HTL, CTL and B cell epitopes of UK variant strain and the remaining 33 epitopes are conserved when compared to other strains subjected to analyses (**Figure S8**). These results suggested that all the epitopes selected for the MEVC had high degree of conservancy among all the variant strains belonging various distinct clades.

### 3.10. Designing of MEVC

The antigenic 14 HTL and 10 CTL epitopes possessing the highest affinity for the respective HLA alleles and four B-cell epitopes that displayed non-allergenic, non-toxic and immunogenic features were selected for incorporation into the multi-epitope vaccine construct. The adjuvant β-defensin was coupled at the N terminal by EAAAK linker with B cell epitope and subsequently, AAY, GPGPG and KK linkers were used to couple B-cell epitopes, CTL epitopes and B HTL epitopes respectively. Adjuvants like Universal memory T cell helper peptide (TpD), PADRE (Pan HLA-DR reactive epitope) and an M cell ligand were coupled by using EAAAK linkers into the vaccine construct. HHHHHH was coupled at C terminal by EAAAK linker for the easy purification of the vaccine (Fig. 1). The final multi-epitope vaccine construct was composed of 575 amino acid residues, which was then validated for antigenic, allergenic and physiochemical properties.

**Figure 1.**
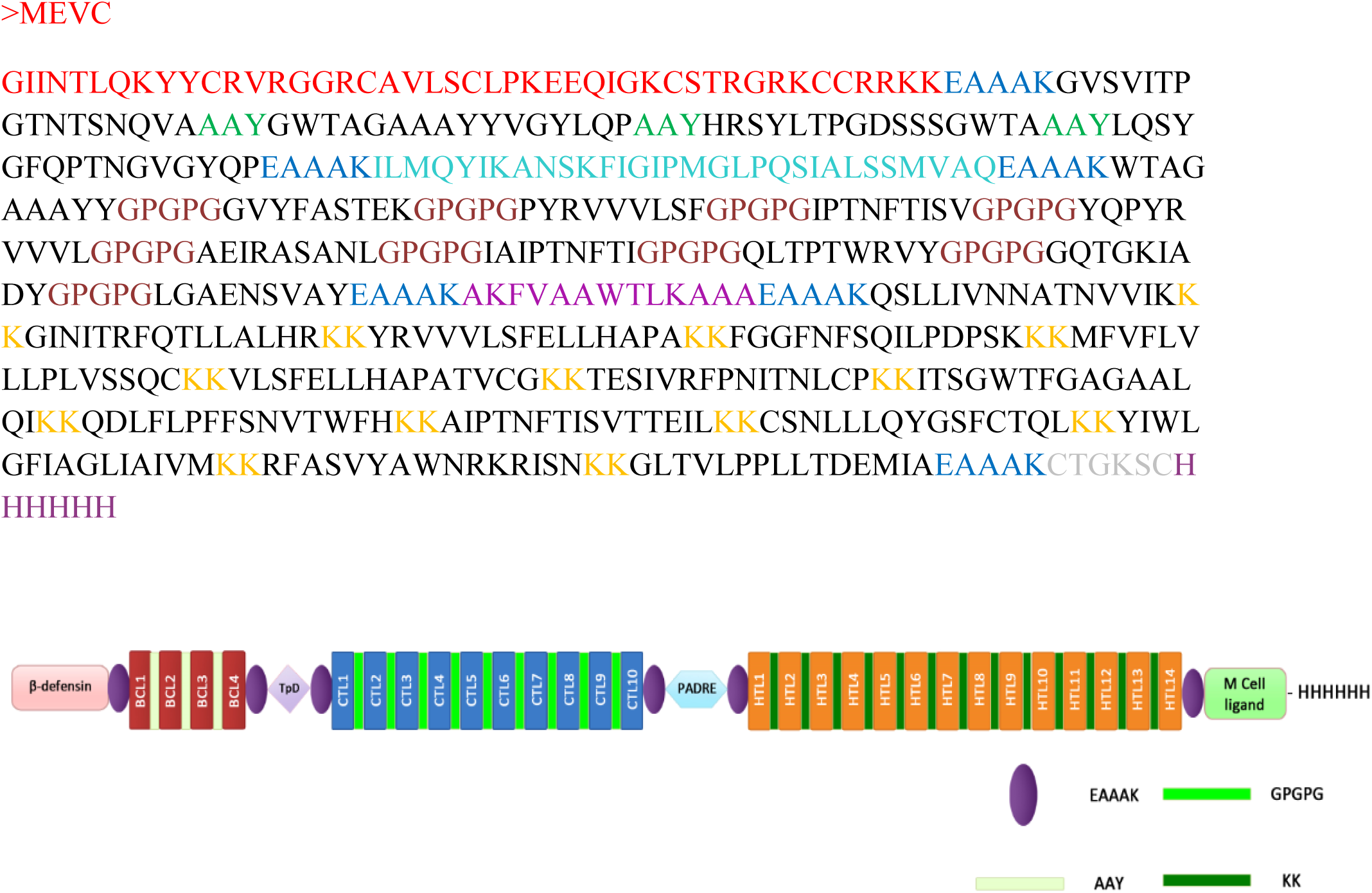
Multi epitope vaccine construct.

#### 3.10.1. Physiochemical properties, antigenicity and allergenicity of MEVC

Analysis of the physiochemical properties of a multi-subunit vaccine helps in analyzing the optimal immune response of the vaccine and its stability inside the host. The physiochemical properties of the multi epitope vaccine construct was calculated by Protparam tool (**Table 4**). The total number of amino acids in the MEVC was 575 with the molecular weight of 61480.71. The total number of positively charged residues (Arg + Lys) and negatively charged residues (Asp + Glu) were 67 and 21 respectively. The theoretical isoelectric point (PI) was calculated as 9.92. The instability index (< 40) indicates that the designed vaccine construct has high stability for the initiation of an immunogenic reaction. The aliphatic index of recombinant protein construct was calculated as 84.07 and this indicates that the construct has stability in various temperatures. Grand Average of Hydropathicity (GRAVY) was 0.001.

Antigenicity is a primary requisite of a successful vaccine candidate. A vaccine construct must possess both immunogenicity and antigenicity to elicit both humoral and cell-mediated immune responses. Upon analyzing the vaccine construct sequence in VaxiJen server, the constructed vaccine was found to be antigenic in nature with an overall prediction score of 0.6077. The score signifies the antigenic potential of the vaccine construct that it can evoke the immune response inside the host. Allergenicity and toxicity analyses revealed the non-allergenic and non-toxic behavior of the vaccine construct. In summary, the constructed epitope was observed to be stable, soluble, antigenic, non-allergenic and nontoxic.

#### 3.10.2. Secondary and Tertiary structure prediction and validation

Secondary and tertiary structure prediction helps in functional annotation of the multi-epitope vaccines. It also helps in analyzing the interaction of this vaccine construct with the immunological receptors like TLRs. Secondary structure of the construct was analyzed by using SOPMA server that revealed the presence of ∼29.57 % α-helix, ∼24.35 % β-sheet, ∼37.91 % coils and ∼8.17% β-turns in the vaccine construct (Fig. 2).

**Figure 2.**
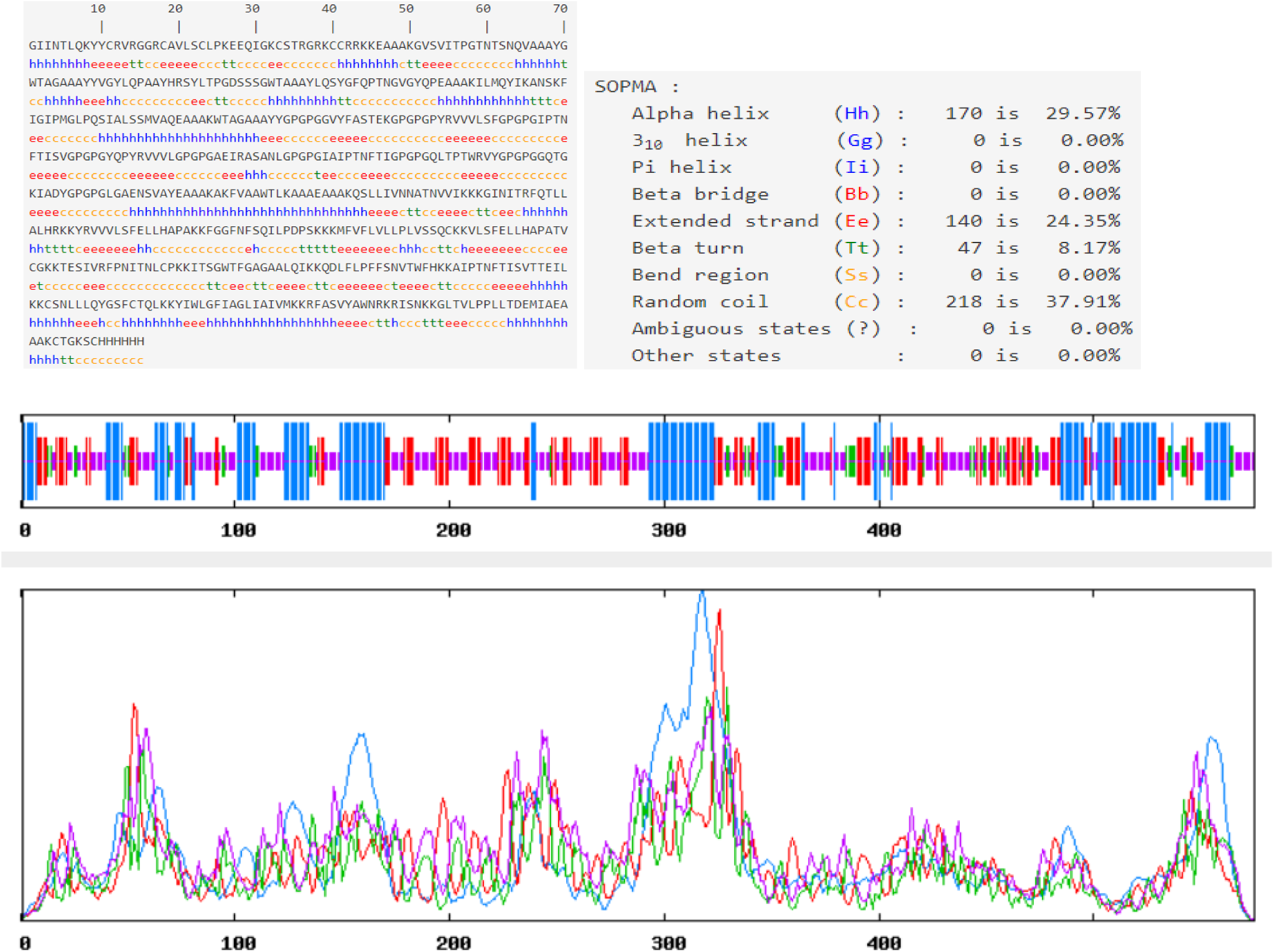
Secondary structure prediction of MEVC.

Tertiary structure of the MEVC was predicted by Galaxy WEB server and refined by Galaxy Refine (Fig. 3a) that performed repeated structure perturbation and subsequent overall structural relaxation by molecular dynamics simulation. For the selected best model, the GDT-HA, RMSD and MolProbity scores were −0.9962, −0.243 and 2.279 respectively. PROCHECK was used to check the stereochemical quality of the structure by considering residue geometry and overall structural geometry. Ramachandran plot analysis of the modelled structure revealed the presence of 89.8% residues in the most favored regions and 8.2% in the additionally allowed regions, 1.2% in the generously allowed region, and 0.8% in the disallowed region (Fig. 3b). To check the potential errors of the protein 3D model, ProSA was used, and it predicted the negative Z-score of −3.65 suggesting good quality of the model (Fig. 3c). These results substantiated the quality of the predicted model.

**Figure 3.**
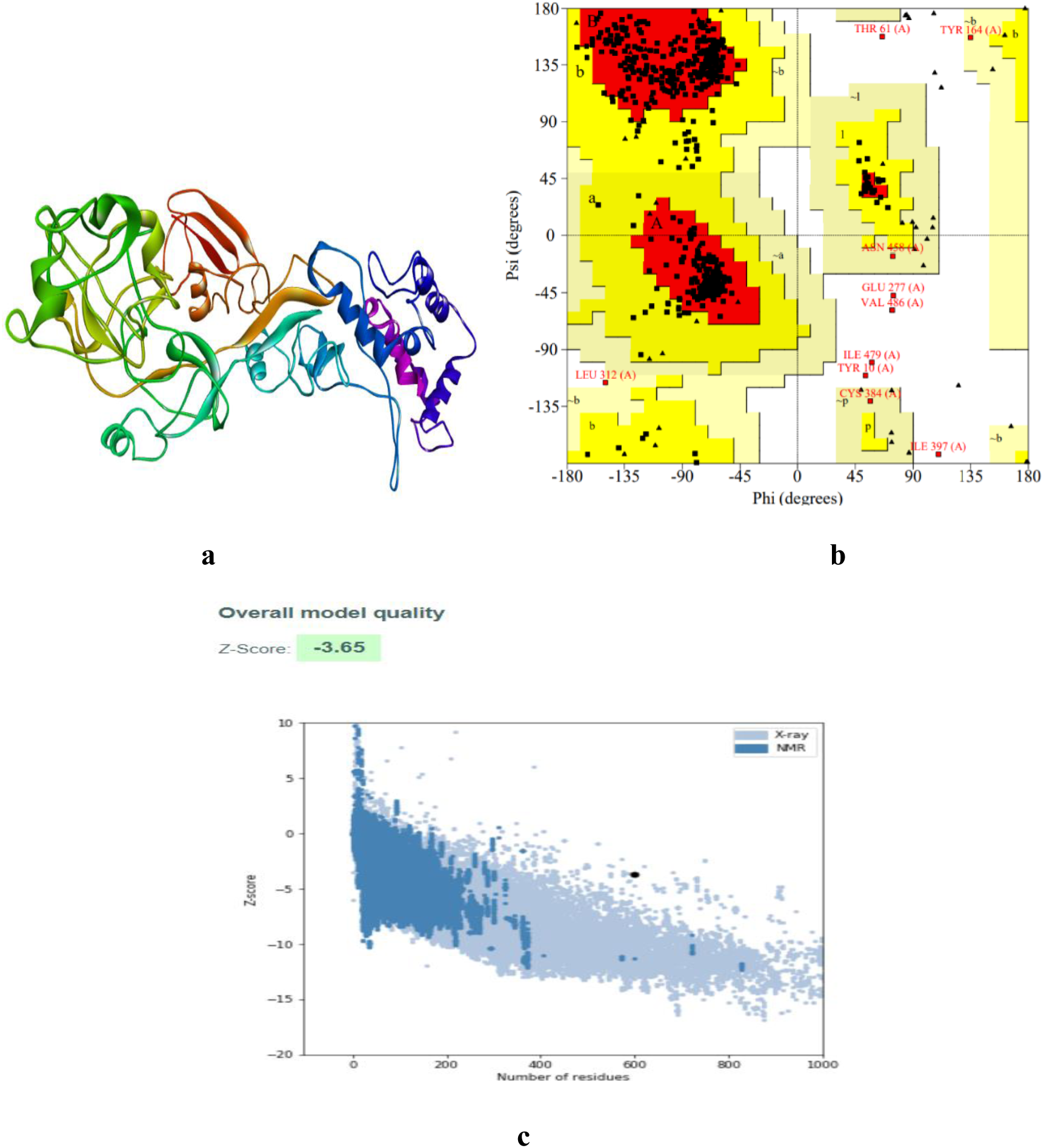
Tertiary structure prediction of MEVC.

#### 3.10.3. Docking of MEVC with receptors

The 3D structures of human TLR3 and TLR4 were retrieved from protein data bank (PDB ID: 2A0Z and 3FXI). Molecular docking analysis was performed using the Cluspro v.2 protein-protein docking server to analyze the interaction pattern of vaccine with TLR3 and TLR4. The server provides cluster scores based on rigid docking by sampling billions of conformation, pairwise RMSD energy minimization. Cluspro v.2 predicted 29 models each of vaccine receptor TLR3 complex and TLR4 complex with their corresponding cluster scores **(Table S8, S9)**. Among these models, the model number 1 (cluster 0) in TLR3 and TLR4 complex were selected as a best docked complex with the lowest energy score of −1274.5 with 29 members (TLR3) and lowest energy score of −1329.1 with 29 members (TLR4). This signifies potential molecular interaction between predicted vaccine construct with TLR3 and TLR4 receptors **(Fig. S9 and Fig. S10)**.

### 3.11. Population coverage of MEVC

Population coverage gives information about the efficiency of epitopes to generate immune response in different population groups located in geographical locations across the world. This analysis is based on the distribution of HLA alleles among different population groups throughout the world. As vaccine construct of this study harbors HTL epitopes for 14 MHC class II and CTL epitopes for 10 MHC class I alleles, the occurrence of these HLA alleles depicted the population coverage of 99.9% throughout the world. Nine countries showed 100% population coverage while ≥ 99% of population was covered in 35 countries whereas ≥ 95% was covered by 23 countries. In 10 countries, ≥ 90% of population was covered. **(Table S10)**. Hence, the vaccine construct showed 90-100% population coverage in 77 countries and 99.9% population coverage throughout the world (**Fig. S11**).

### 3.12. Immune simulations of vaccine construct

C-ImmSim simulator was used to analyse the immune response produced by the final vaccine construct. The tool utilizes the Celada-Seiden model for describing both humoral and cellular profiles of a mammalian immune system against designed vaccine. The immune system simulation server provides an opportunity to study the overall immunogenicity of the generic protein sequence in the context of its amino acid sequence (Rapin, 2011). The total simulation is focused on three events: 1) B-cell epitopes binding, 2) HLA class I and II epitopes binding, and 3) TCR binding, which HLA-peptide complex interaction should be presented. Such processes are independently conducted by cells described by agents and occupy a specified simulated biological amount. The tool generated the immune response simulations that match the response of a real immune system **(Fig. S12; a-j)**. The cumulative results of immune responses after three times antigen exposure revealed that the primary immune response against the antigenic fragments was elevated and it was indicated by gradual increase of IgM level after each antigen exposure. Similarly, the secondary response was characterized by adequate generation of IgM+IgG more than IgM. An increased level of IgG1+IgG2 and IgG1 (**Fig. S12a**) was also observed. On the subsequent exposure of vaccine, a decrease in the level of antigens was observed indicating the development of immunogenic response in the form of immune memory. The elevated levels of all circulating immunoglobulins indicate the accuracy of relevant clonal proliferation of B-cell and T-cell population. Furthermore, an increase in the B-cell population (**Fig. S12; b and c**) was characterized by an increase in the expression of immunoglobulins, which resulted in a decrease in the concentration of the antigen. Besides, there was a consistent rise in Th (helper) and Tc (cytotoxic) cell population with memory development (**Fig. S12; d, e, f**). Total NK cells, dendritic cells and macrophages were also increased (**Fig. S12; g, h, i**). It was also observed that the production of IFN-gamma was stimulated after immunization (**Fig. S12j**). These results revealed that the MEVC proposed in this study could generate a strong immune response, and immunity increases even on subsequent repeated exposure.

## Discussion

For developing efficient epitope-based peptide vaccine, surface glycoprotein of virus is considered as the major focus by vaccine design platforms as these proteins are involved in the interaction between cell receptor and virus particle and play a crucial role in the pathogenesis of the disease (Tilston-Lunel *et al*., 2016). In the present study, we attempted reverse vaccinology approach for designing of a multi-epitope vaccine based on the Spike (S) protein of SARS-CoV-2 that may efficiently elicit humoral and cellular mediated immune responses against the viral infection. Substitution mutations in spike protein sequences of SARS-CoV-2 isolates from COVID-19 positive clinical samples of Tamilnadu, India, 10 other strains representing various clades and recently originated UK variant strains were included in the analysis in comparison with the reference sequence of Wuhan strains and a phylogenetic tree was constructed employing MEGA. Among the 31 sequences, a sequence of an isolate was selected to predict various B-cell and T-cell epitopes against SARS-CoV-2.

The retrieved structural protein and its antigenicity score suggests that the spike protein is the most potent protein to generate immune response. Both T and B cell epitopes were predicted using the immunoinformatics tools. MHC class I binding peptides generally have 8-11 amino acids while MHC class II binding peptides are typically 12-25 amino acids long. The study results on T cell epitope prediction and analyses based on features such as antigenicity, allergenicity, immunogenicity, conservancy and toxicity suggested that the selected T cell epitopes were having high scores for these features. The lower percentile rank and lower IC50 (<100nM for Class I and <1000 nM for Class II) values for alleles of T cell epitopes met the criteria to be strong binders supporting the allele selection for population coverage. Further, the B-cell epitopes were divided into two main categories such as continuous or linear B-cell epitope and discontinuous or conformational B-cell epitope. The predicted linear B cell epitopes with higher cutoff values (0.9 and above) were analyzed for antigenicity, allergenicity, toxicity and conservancy and the best scoring epitopes were selected for multi epitope vaccine construct. The secondary and tertiary structures of the spike protein sequence were predicted and validated to identify conformational or discontinuous B cell epitopes. Totally, 6 epitopes were predicted with high score.

Vaccine construct should be antigenic, non-allergenic and non-toxic to make it a potent vaccine candidate against SARS-CoV-2. Hence, 28 epitopes such as 10 CTL, 14 HTL and 12 B cell epitopes were chosen for MEVC design from the promising 36 epitopes selected for the study based on conservancy, antigenicity, nontoxicity and non-allergenicity features. The two B cell epitopes were not chosen due to toxicity and allergenicity features to the human. So, the selected T and B cell epitopes were incorporated in the multi epitope vaccine construct. It is noteworthy that the 36 epitopes selected for the MEVC were having high degree of conservancy among all the variant strains belonging various distinct clades including the recently emerged UK variant SARS-CoV-2 which had only 3 mutations in 3 epitopes (each in one of the HTL, CTL and B cell epitopes) out of the total 36 epitopes. Hence, the epitopes used in the MEVC are highly conserved against SARS-CoV-2 variants of various clades.

These epitopes were linked by adjuvants and linker molecules. Four different adjuvants such as β-defensin, TpD, PADRE and M cell ligand were added to the MEVC to enhance the innate and adaptive immune responses besides aiding the transportation of MEVC through the intestinal membrane barrier. Adjuvants are the small peptides that when added along with the antigenic vaccine can enhance the specific immunogenic response by influencing the onset, strength, and longevity to antigens and induce protection against infection (Lee and Nguyen, 2015; Schijns and Lavelle, 2011). The used linkers included GPGPG (Nezafat *et al*., 2014), AAY (Schubert and Kohlbacher, 2016), KK (Nezafat *et al*., 2017) and they were chosen based on the linkers for the design of multi-epitope peptide vaccines in literature and linker databases. Analysis of the physiochemical properties of 575 residues length MVEC revealed that instability index (26.87) classified the protein as stable and the aliphatic index (84.07) indicated that this protein could have stability in several temperatures. This construct showed high antigenicity and it was non-allergic and non-toxic. The secondary and tertiary structure predictions revealed the structural details of the vaccine construct. The modelled 3D structure was further refined that increased its overall quality. Protein-protein docking was performed to know the interaction of vaccine construct and immune receptors. Various immune receptors like Toll-like receptors (TLRs) act as sensor proteins that help in recognizing and differentiating self and non-self molecules by recognizing molecular patterns on the surface of infectious agents and elicit immune response against these agents. In this study, the multi epitope vaccine construct was docked with TLR3 and TLR4. The molecular interaction of vaccine with TLR3 and TLR4 through docking analysis suggested that the constructed vaccine possessed a significant affinity towards the toll-like receptors to recognize molecular patterns of pathogen to initiate immune response. Thus, the vaccine TLR complex is capable of generating an effective innate immune response against SARS-CoV-2. The biochemical integrity in epitope structures was further observed by Ramachandran plot analysis.

Based on the presence of epitopes for the HLA alleles, the population coverage of the vaccine showed 99.9% of world population. The results revealed that the selected epitopes and their alleles have ideal population coverage in different geographic regions around the world. Few countries showed 100% coverage and most of the countries showed ≥ 99% population coverage. In total, vaccine construct showed 90-100% population coverage in 77 countries. The consistent increase of high level of IFN gamma supported the activation of humoral immunity. Immunogenic potential of vaccine was understood by repeated administration in 3 injections and the result suggested the consistent generation of a strong immune response. Further, CTL, HTL and B cell epitopes incorporated in the MEVC were analysed with the respective positions of S protein of 3156 downloaded sequences revealed that all these epitopes were highly conserved and these epitopes showed high degree of homology in the range of 97.53% to 100%. With all these immunoinformatic approaches, the vaccine designed against SARS-CoV-2 showed promising results in inducing immune response.

Though the study analyses were made based on spike protein of SARS-CoV-2, an earlier study reported the analyses of CTL, HTL and B cell epitopes from 3 different proteins of the virus such as spike protein (S), membrane glycoprotein (M) and nucleocapsid (N) protein (Kalita *et al*., 2020). The spike protein is a key target for the development of vaccines, therapeutic antibodies, and diagnostics for coronavirus, and hence, it was chosen for the study. Inclusion of more potential epitopes from other proteins in the MEVC may suffer the limitation of complexity of the construct besides the challenges associated with the synthesis. A recent study focused on the single protein (spike protein) to generate multiple epitopes such as 13 for MHC I and 3 for MHC II epitopes (Bhattacharya *et al*., 2020); but, this study has the limitation of not considering the B cell epitopes. Another recent study reported the design of subunit vaccines against SARS-CoV-2 that used only CTL epitopes without considering the significance of B-cell or HTL epitopes (Seema, 2020). Some studies have used other proteins of the virus and restrict the analyses to one of these three epitopes (Joshi *et al*., 2020). The present study has an advantage in providing a potential MEVC as it contains all three types of epitopes such as CTL, HTL and B cell epitopes. This *in silico* study is an attempt to describe the potential immunogenic target over the structural proteins and to propose a novel MVEC, for providing new rays of hope in the initial phase of vaccine development and subsequent experimental validation to confer protection against SARS-CoV-2 infection.

## Acknowledgement

The authors thank Department of Health Research (DHR), Govt of India to the State VRDL, Department of Virology, King Institute of Preventive Medicine and Research, Chennai for the financial support to the lab wherein the study is carried out.

## Supplementary Files

i. **(i) Supplementary Figures**
ii. **(ii) Supplementary Tables**
iii. **(iii) Supplementary File A: SARS-CoV-2 genome sequences for retrieval of spike protein for analyses**
iv. **(iv) Supplementary File B: Mutations in spike protein epitopes of B and T cells**

