## Supplementary material for "Immunoinformatic Approach for the identification of T Cell and B Cell Epitopes in the Surface Glycoprotein and Designing a Potent Multiepitope Vaccine Construct Against SARS-CoV-2 including the new UK variant": epitope docking images.pdf

Docking of CTL epitopes

Epitope1

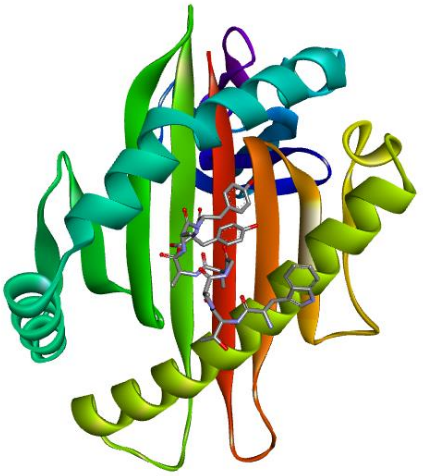

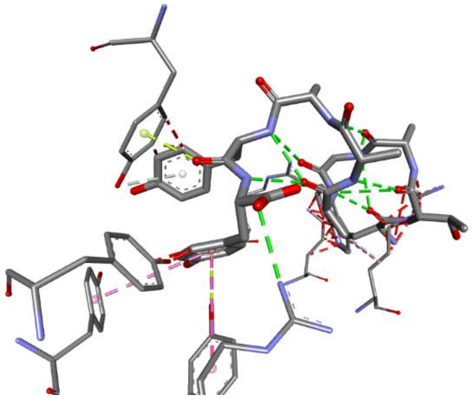

Epitope2

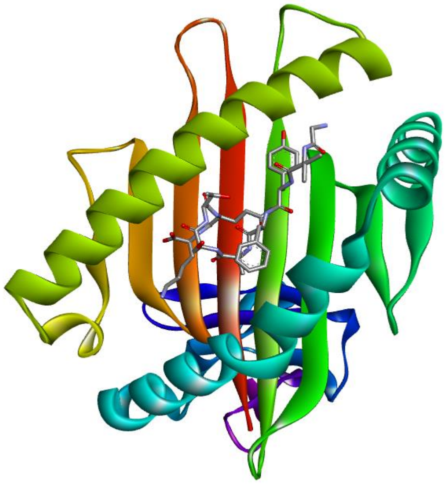

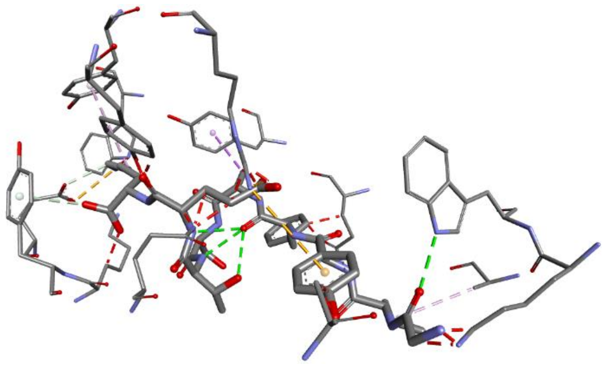

Epitope 3

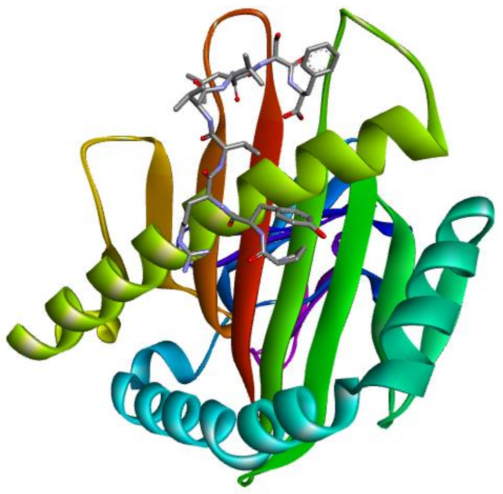

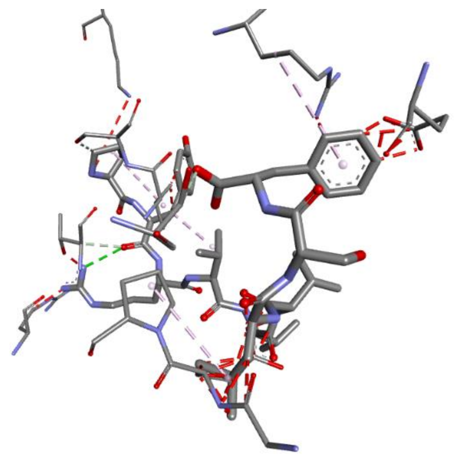

Epitope4

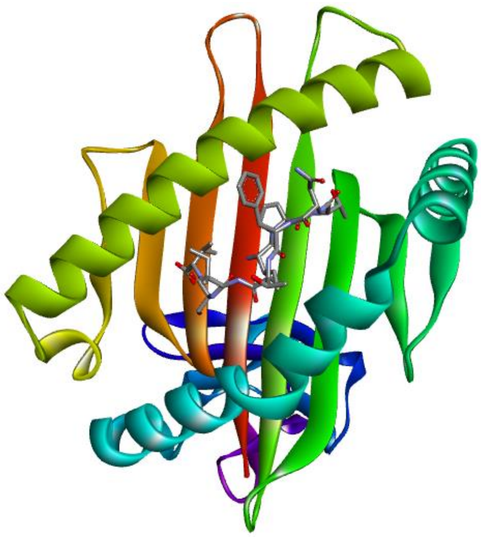

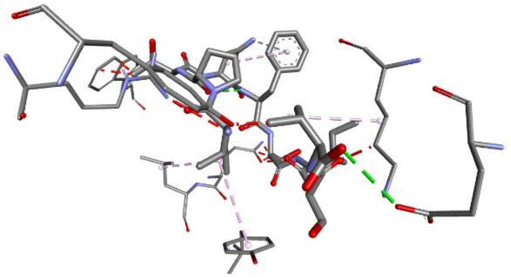

Epitope5

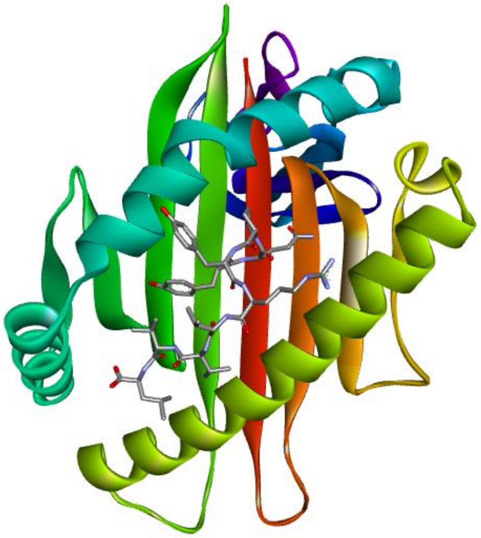

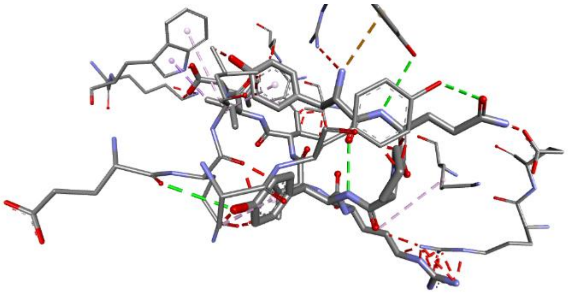

Epitope6

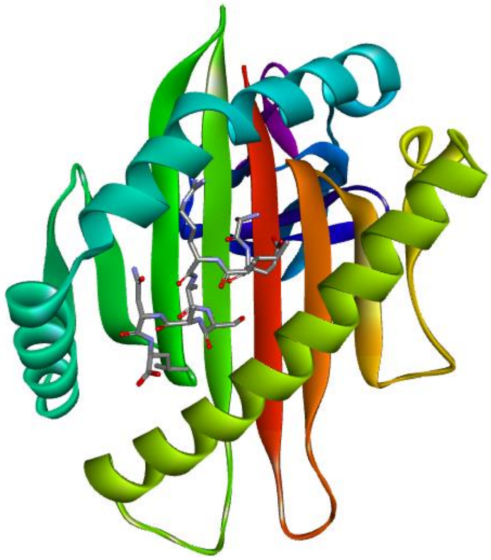

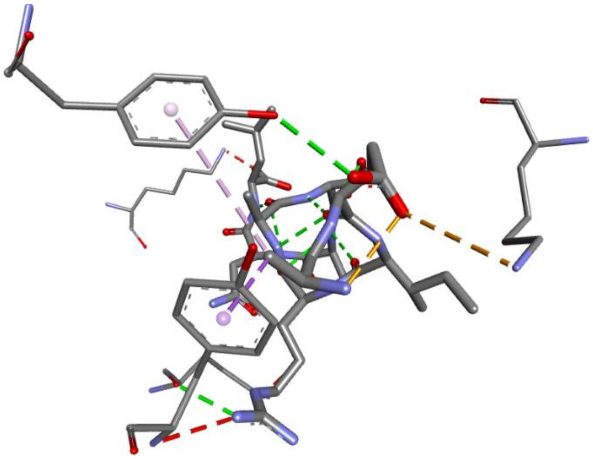

Epitope7

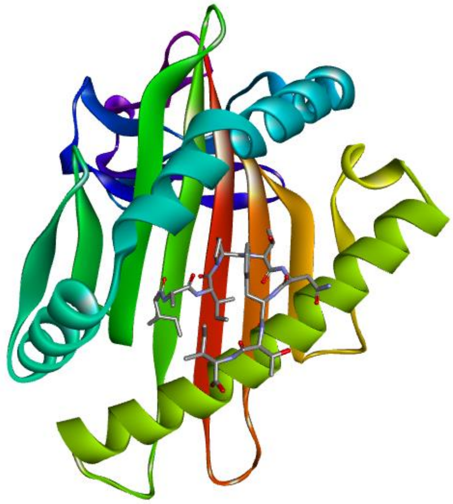

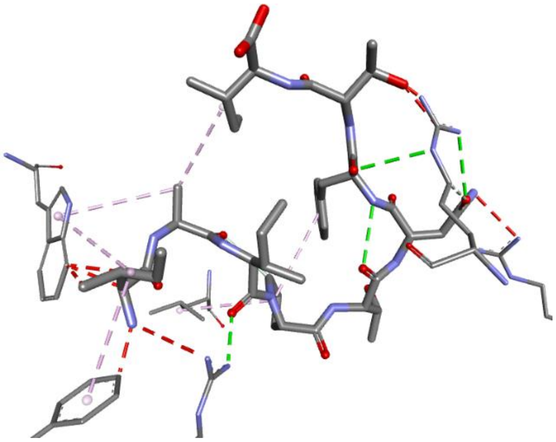

Epitope8

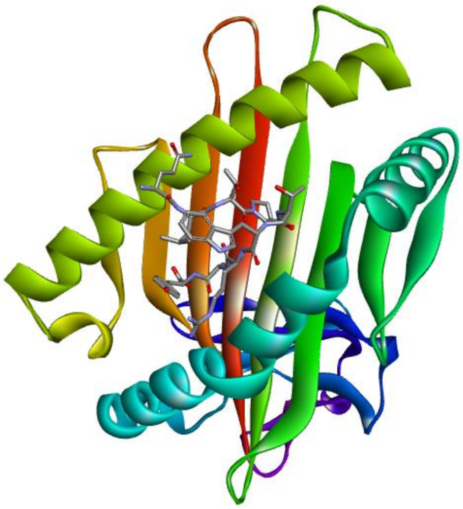

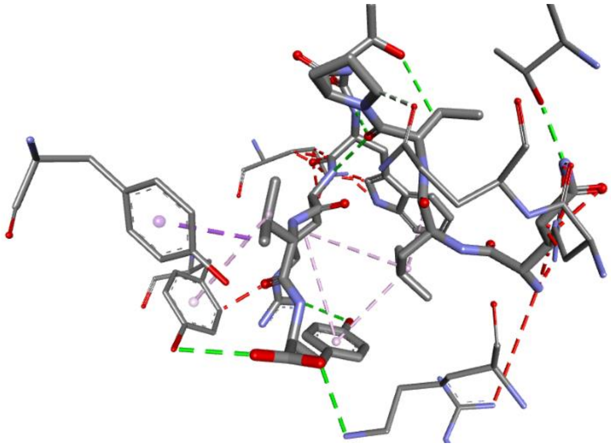

Epitope9

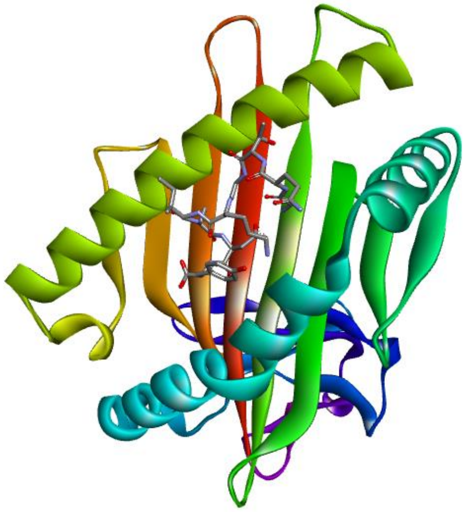

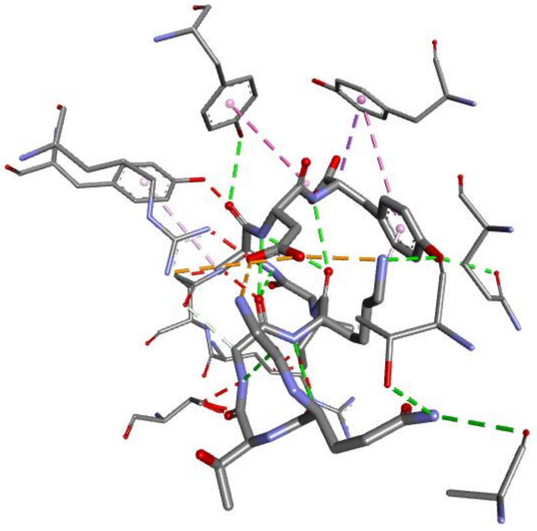

Epitope10

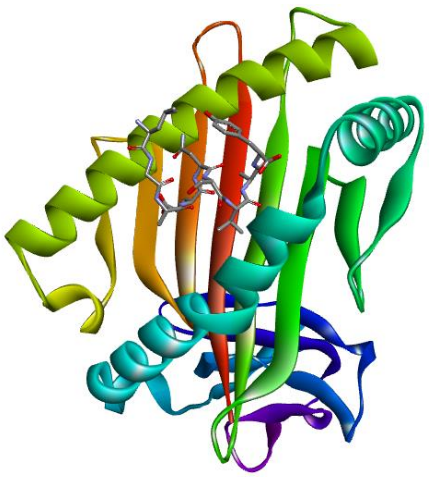

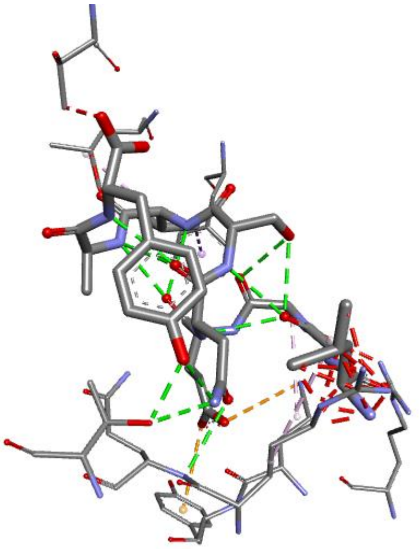

**Class2 MHC**

**EPITOPE1**

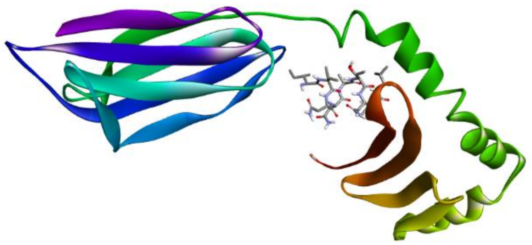

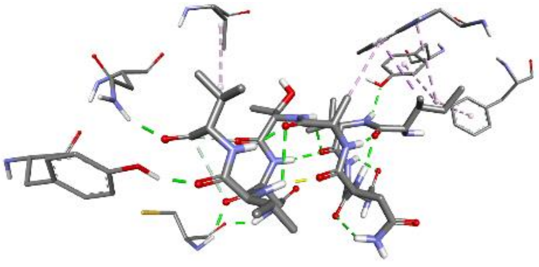

EPITOPE 2

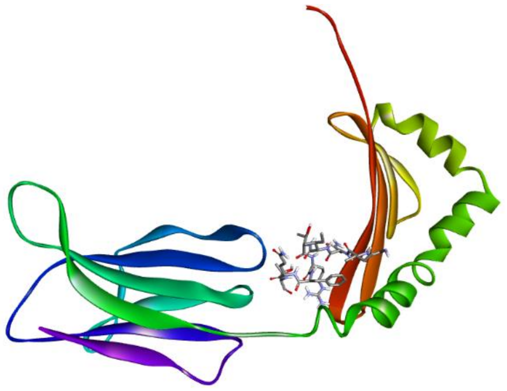

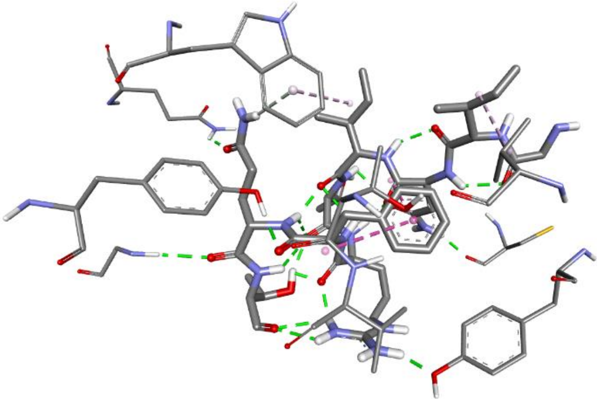

EPITOPE 3

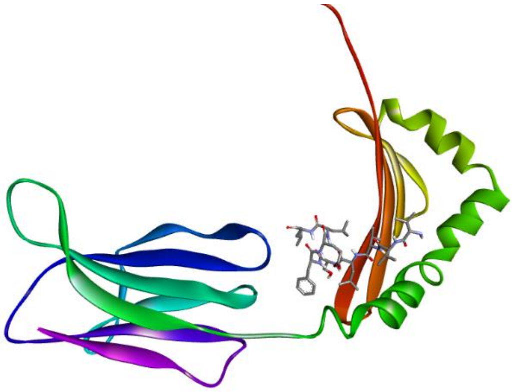

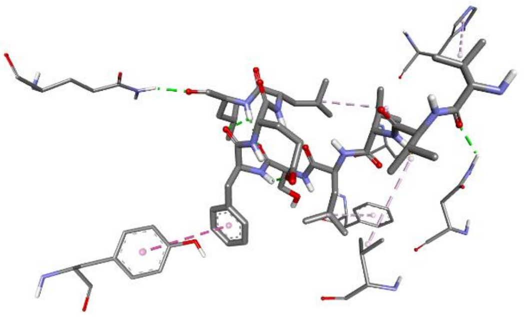

EPITOPE 4

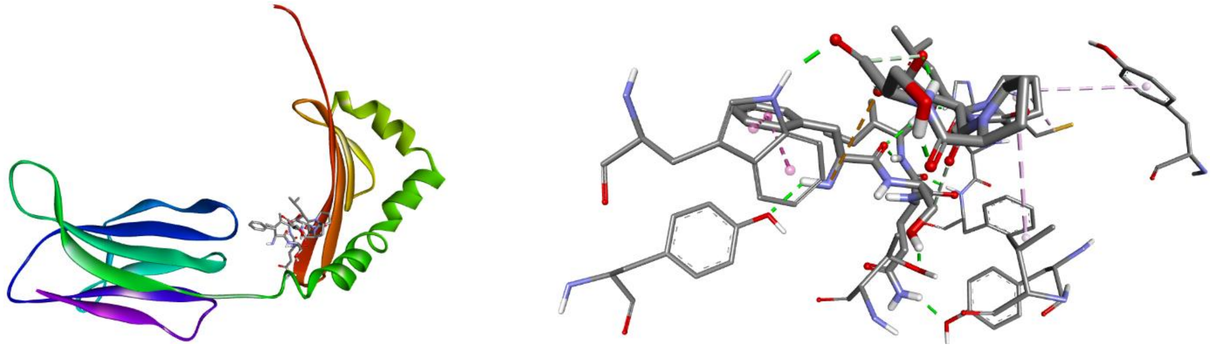

EPITOPE 5

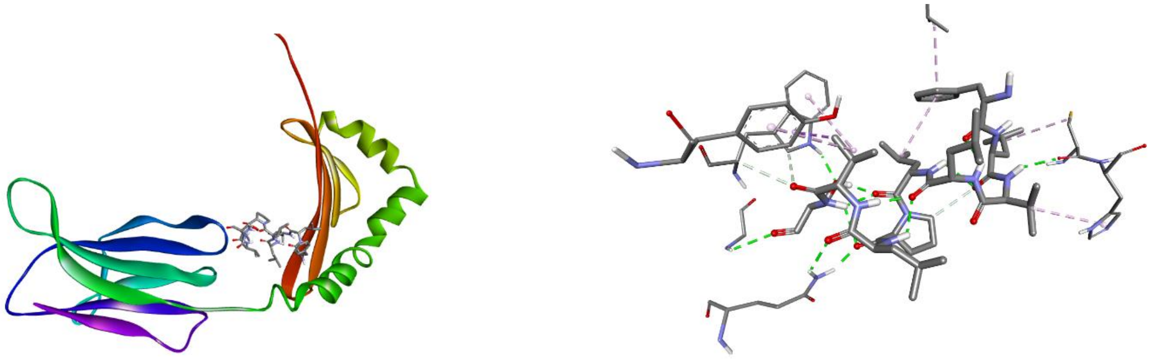

EPITOPE 6

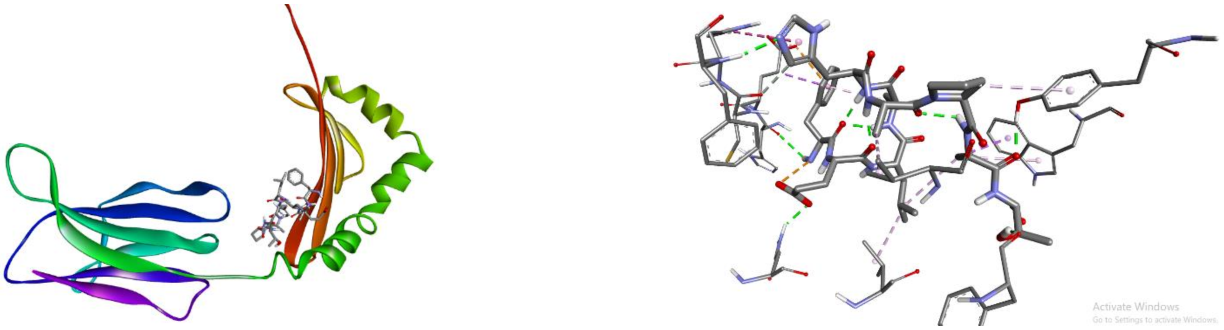

EPITOPE 7

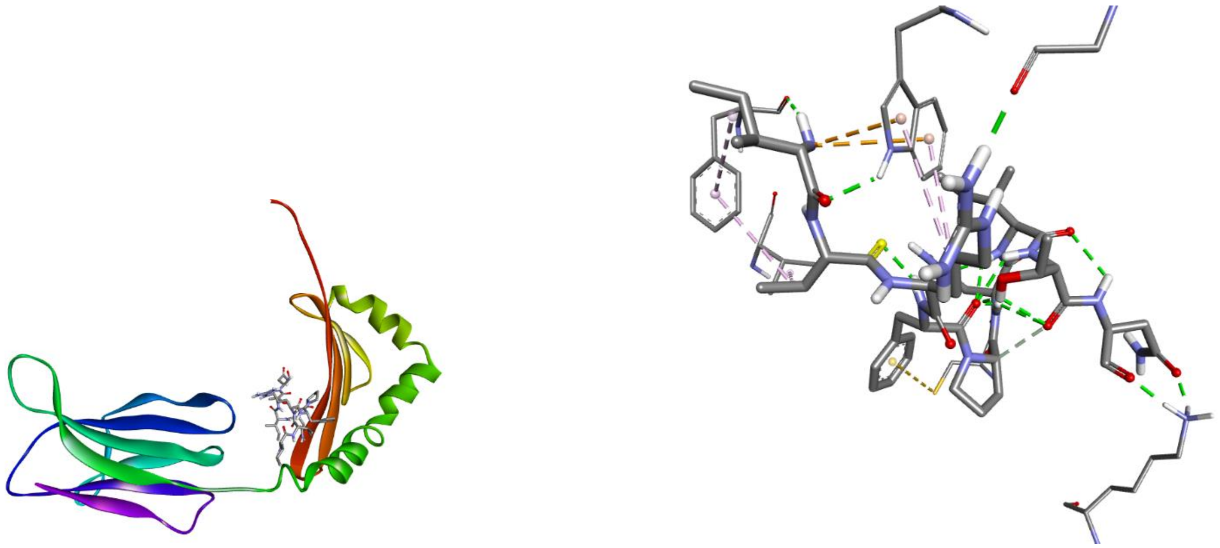

EPITOPE 8

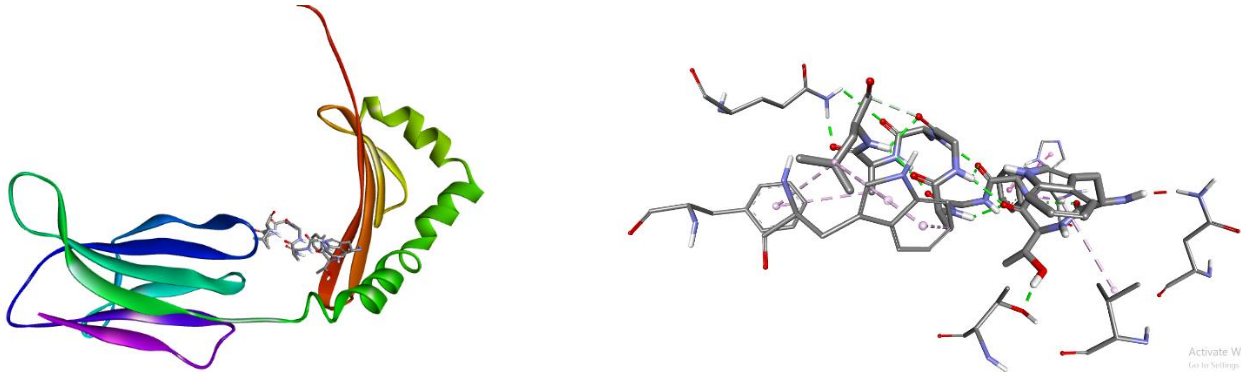

EPITOPE 9

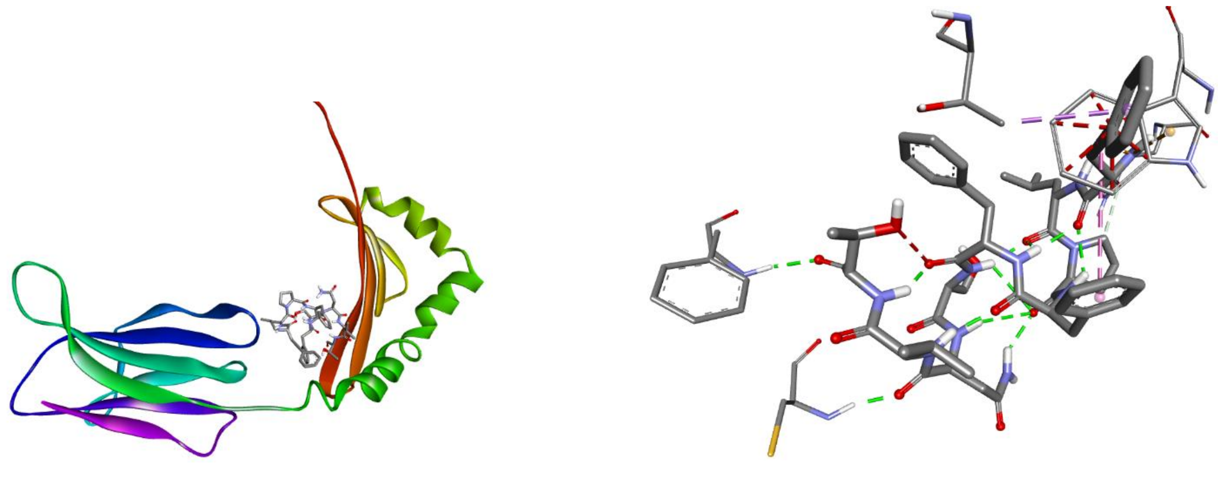

EPITOPE 10

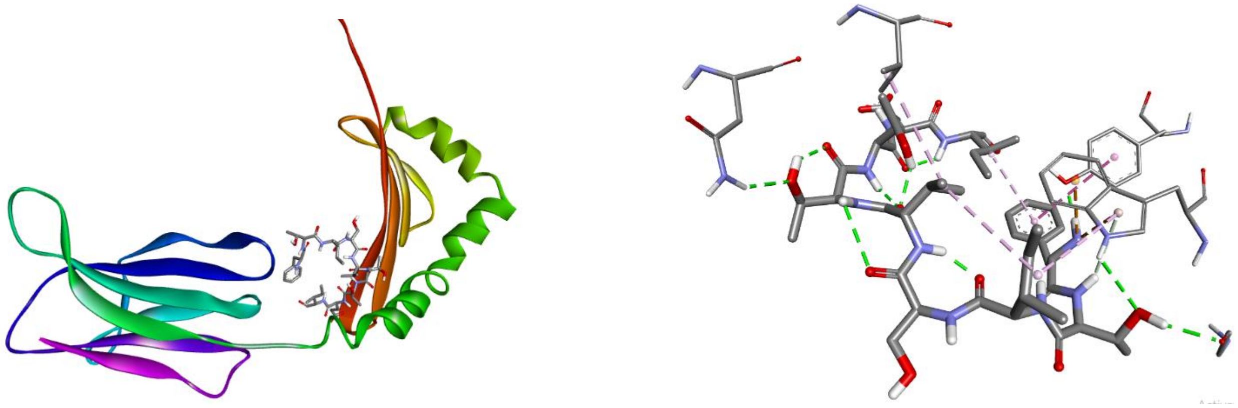

EPITOPE 11

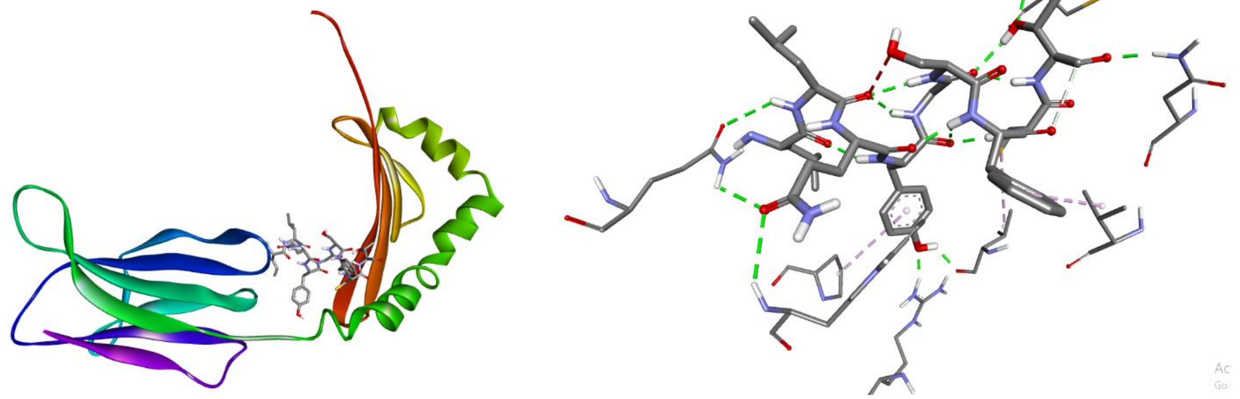

EPITOPE 12

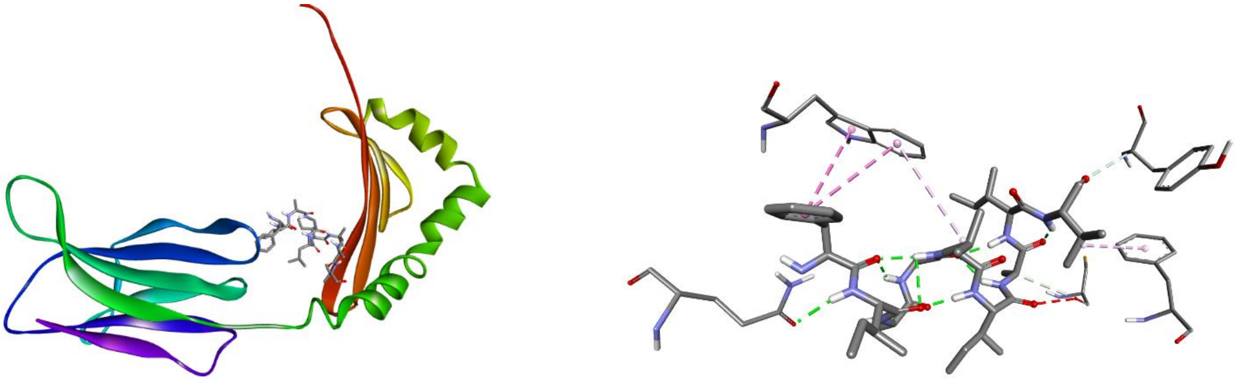

EPITOPE 13

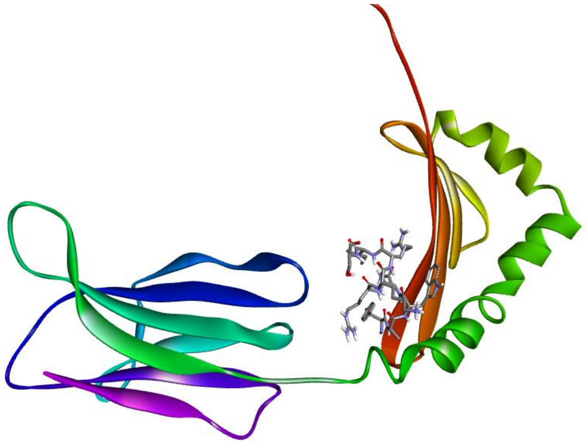 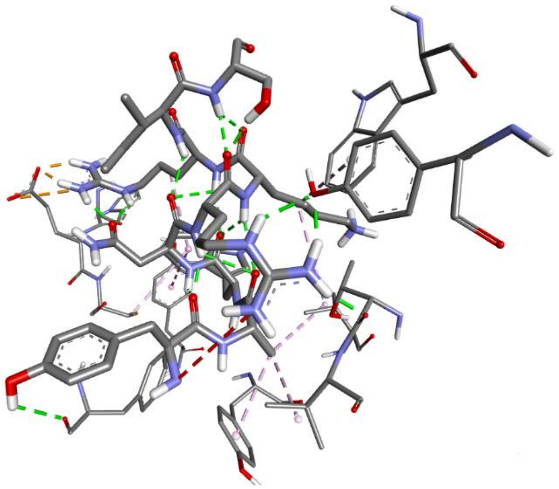

EPITOPE 14

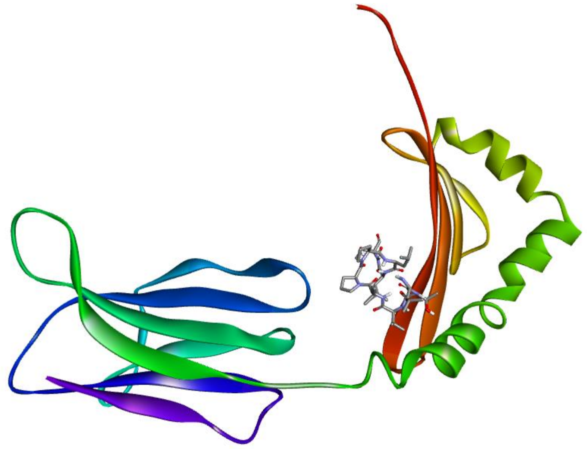 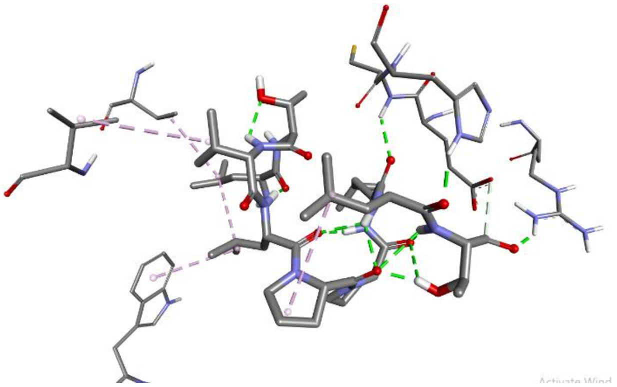

**FOR DISPLAY**

**CTL EPITOPE INTERACTION**

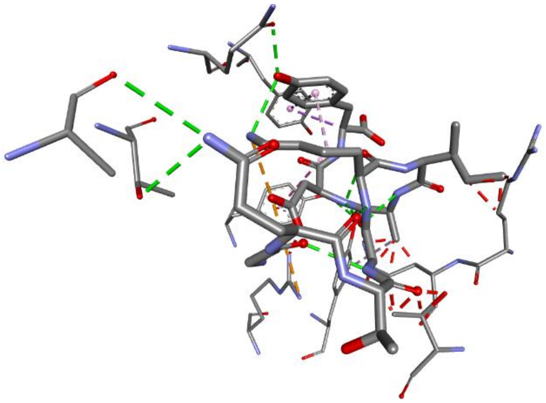

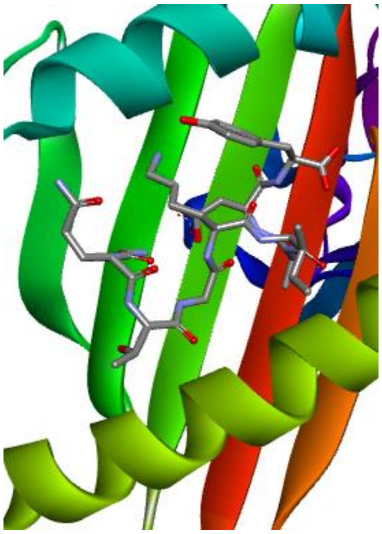

**HTL EPITOPE INTERACTION**

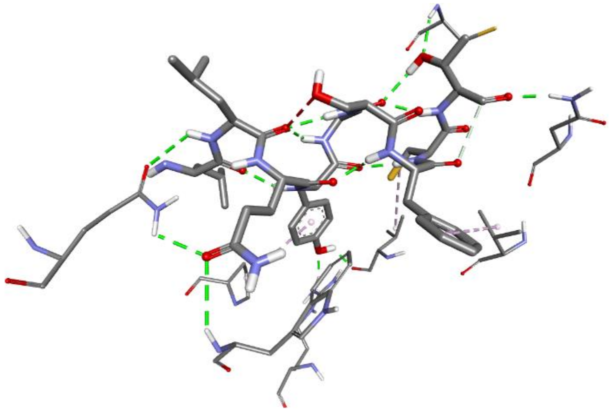

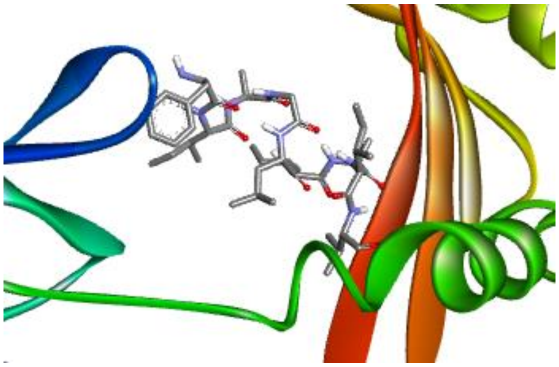
