## Supplementary material for "Immunoinformatic Approach for the identification of T Cell and B Cell Epitopes in the Surface Glycoprotein and Designing a Potent Multiepitope Vaccine Construct Against SARS-CoV-2 including the new UK variant": Supplementary Figures.pdf

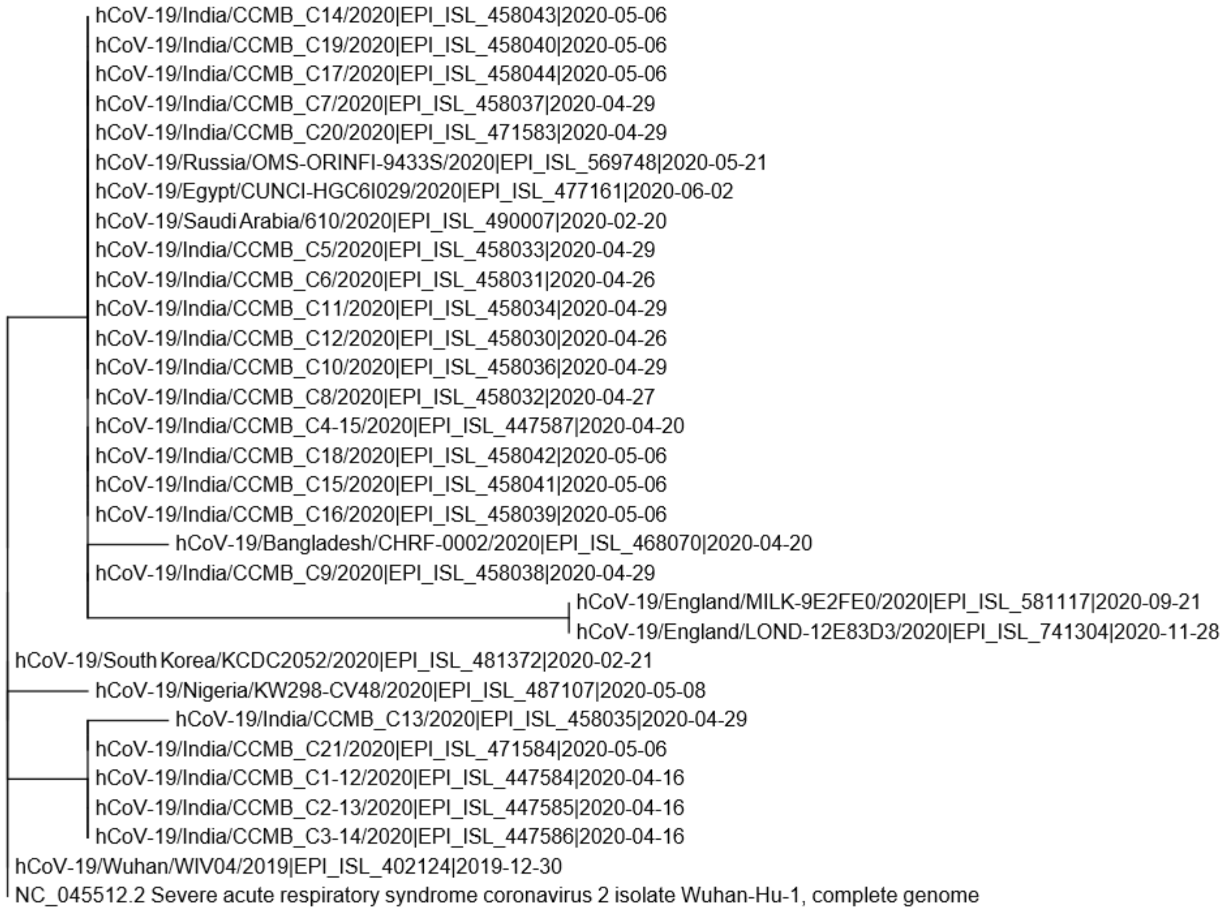


**Fig. S1. Phylogenetic tree constructed from spike protein of SARS-COV-2 isolates.**

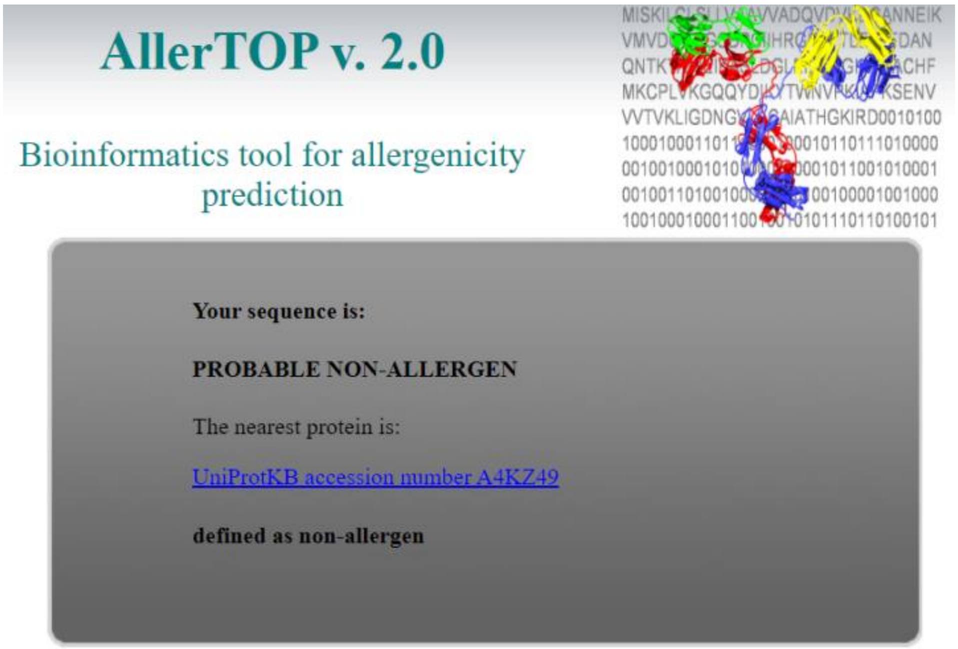


**Fig. S2. AllerTop tool shows that the selected sequence is non-allergic to human.**

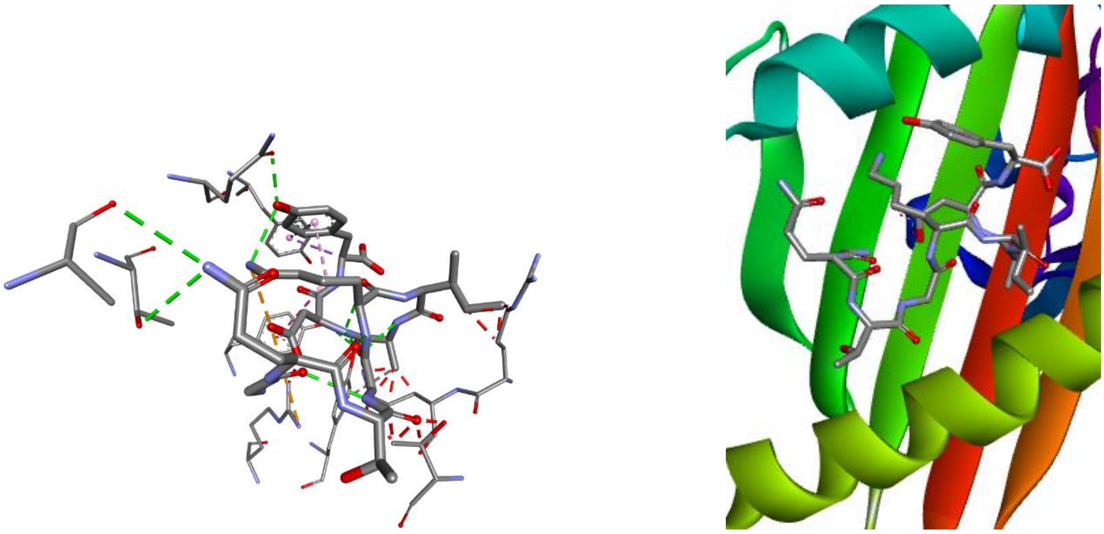

**Fig.S3. CTL epitope interaction**

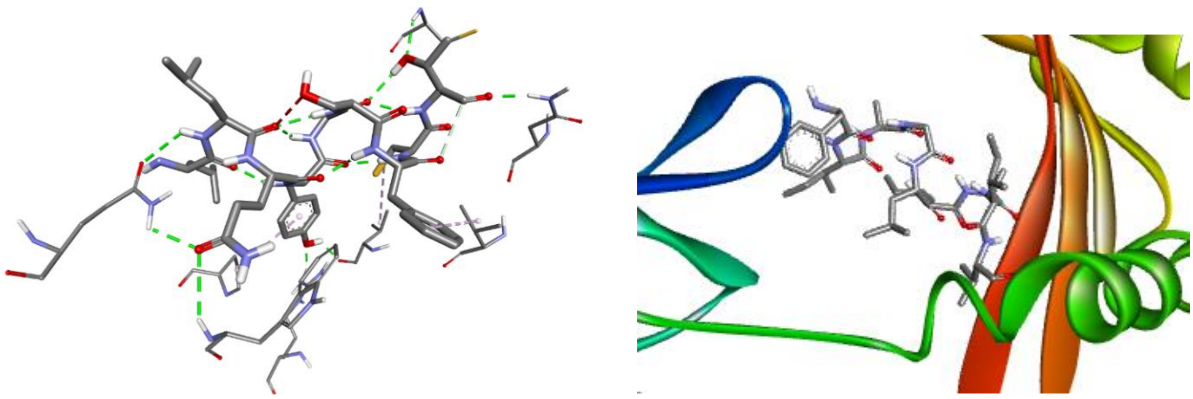

**Fig.S4. HTL epitope interaction**

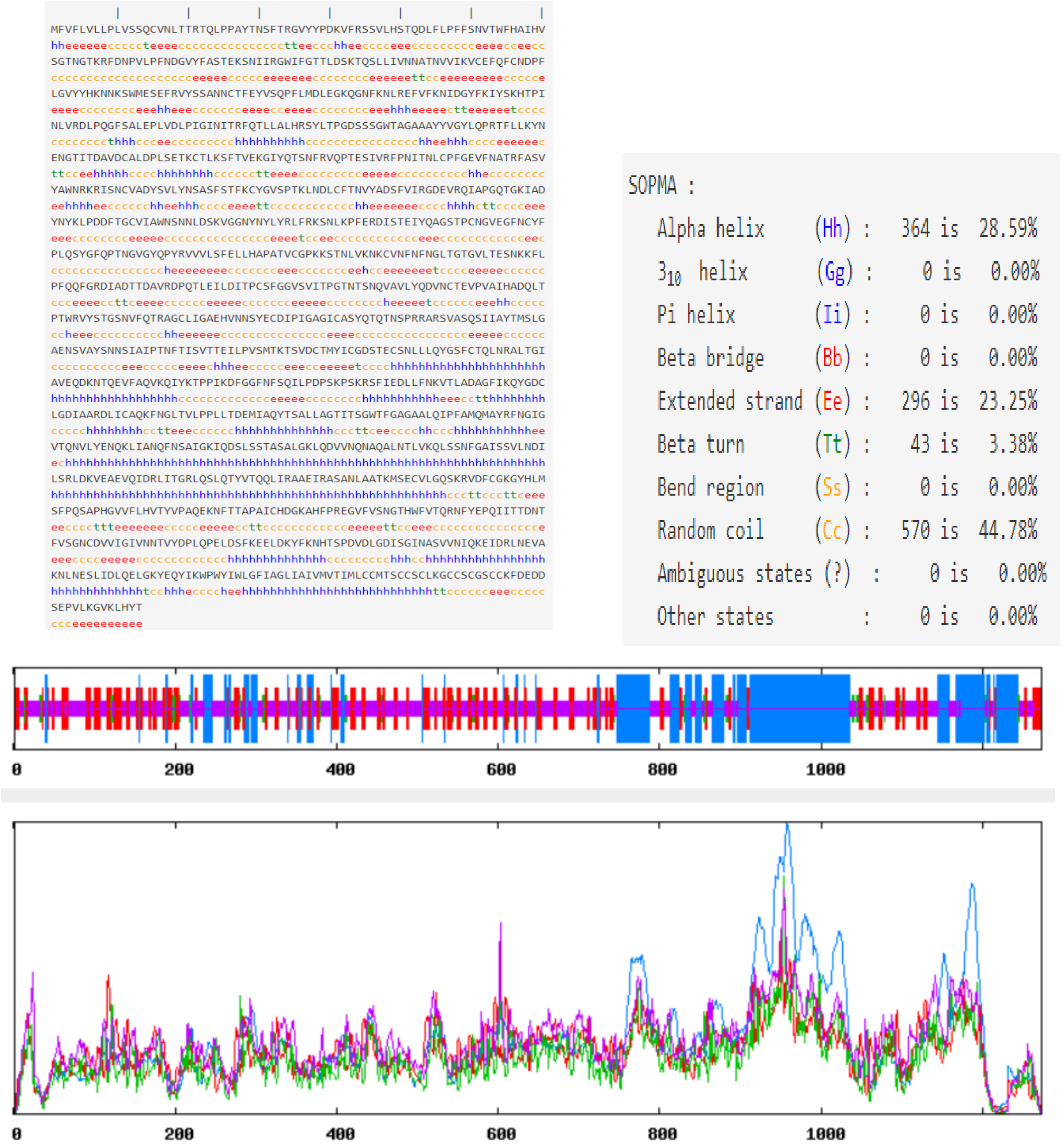


**Fig. S5. Secondary structure prediction of the selected spike sequence**

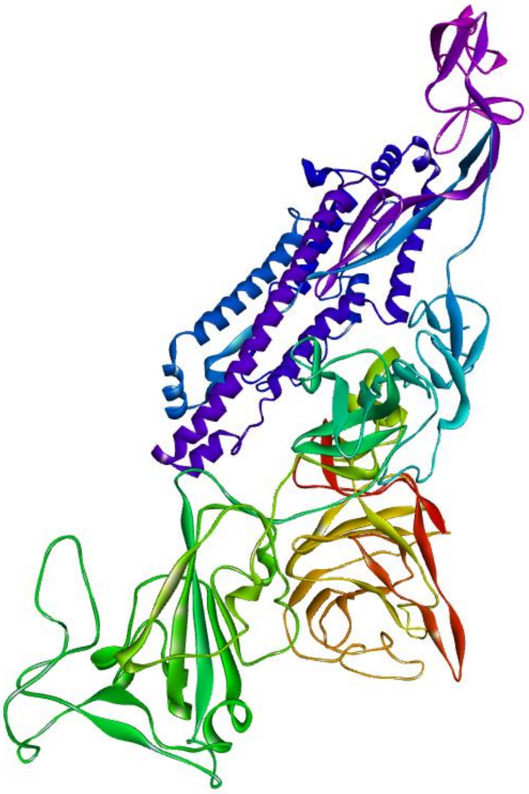

**a**

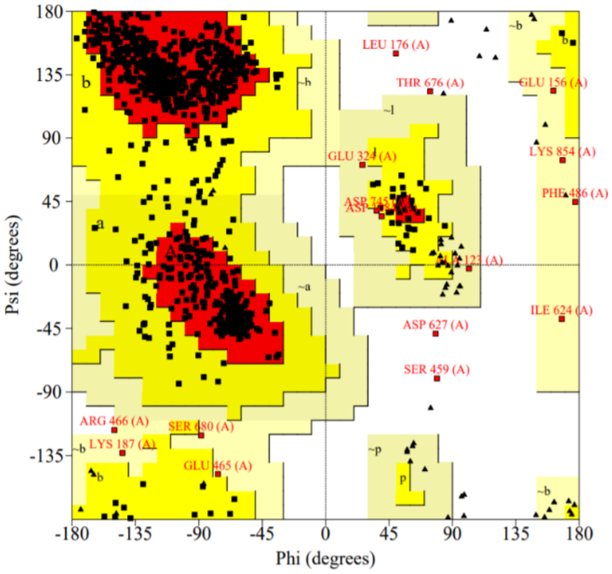


**b**

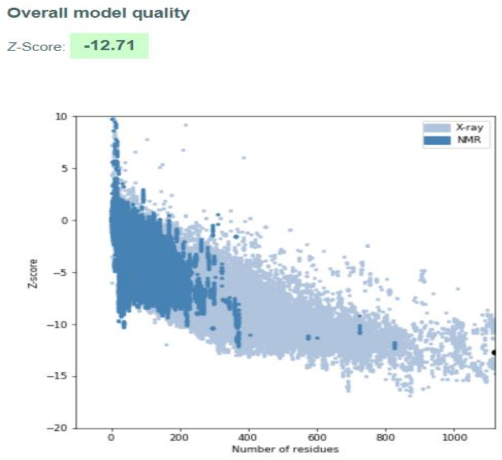


**c**

**Fig. S6. Tertiary structure prediction**

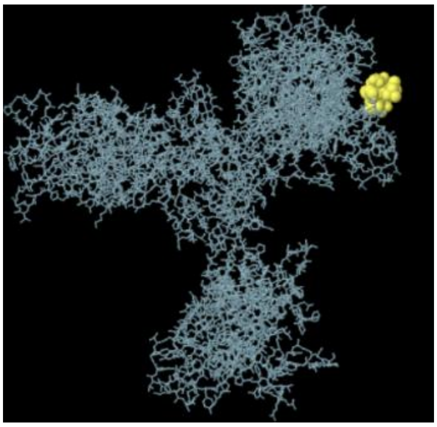

A

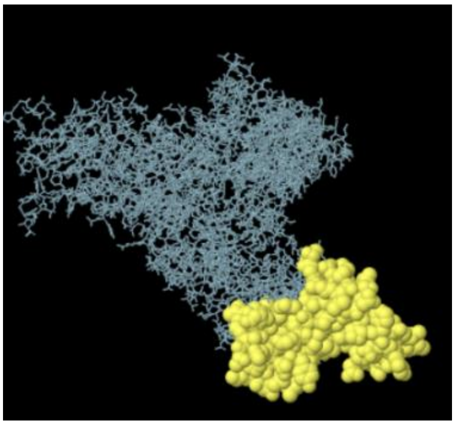

b

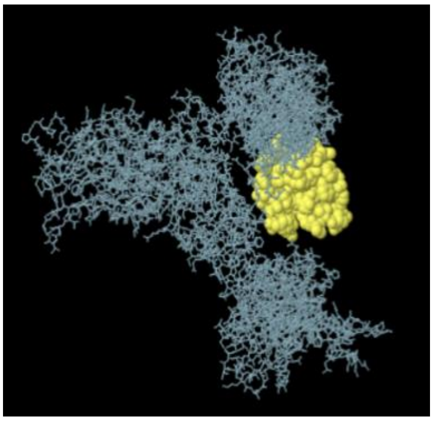

c

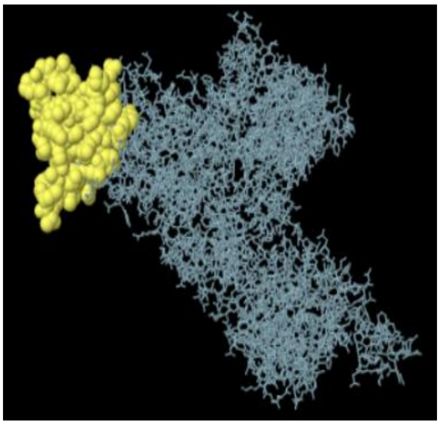

d

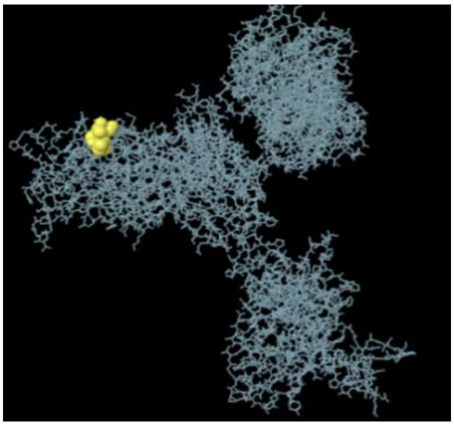

e

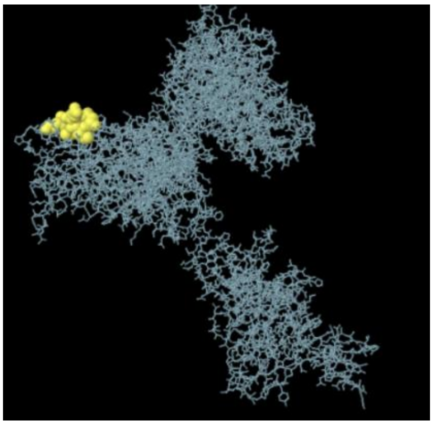

f

**Fig. S7 a-f; Positions of epitopes**

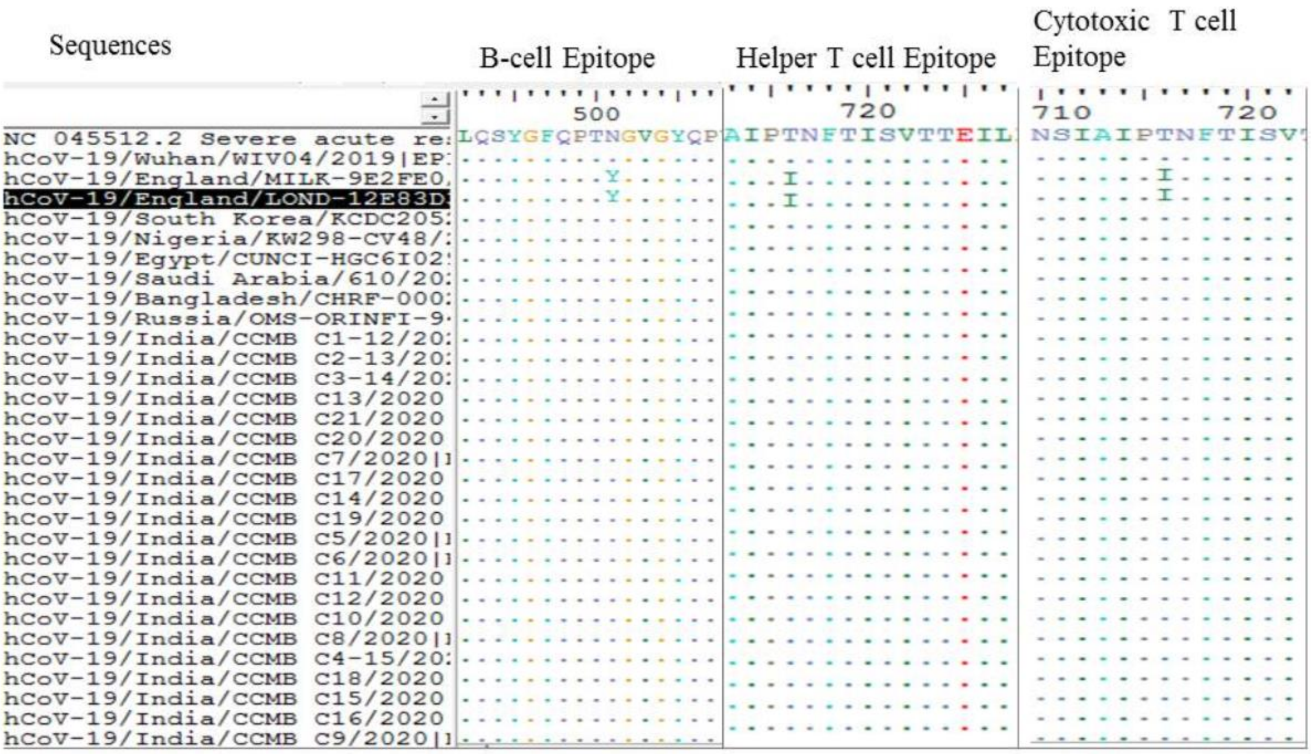


**Fig. S8. Mutations in CTL, HTL and B cell epitopes of new variant UK strain against the other SARS-CoV-2 variants.**

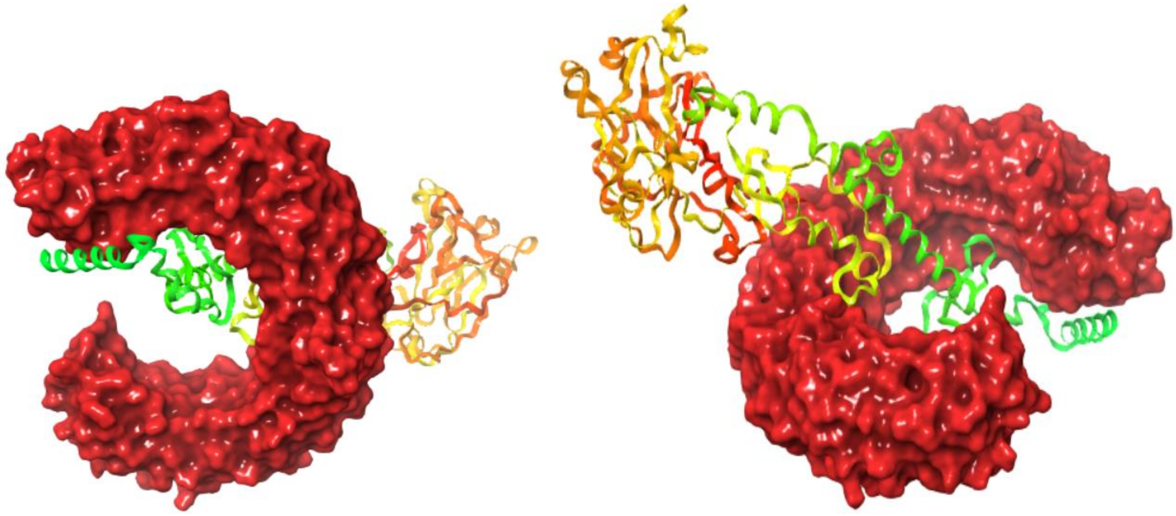

**Fig. S9. Docking with TLR3**

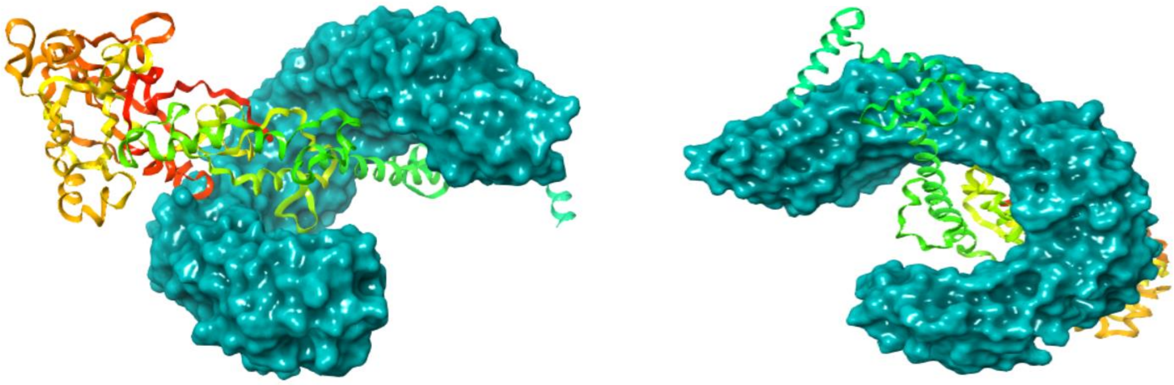

**Fig. S10. Docking with TLR4**

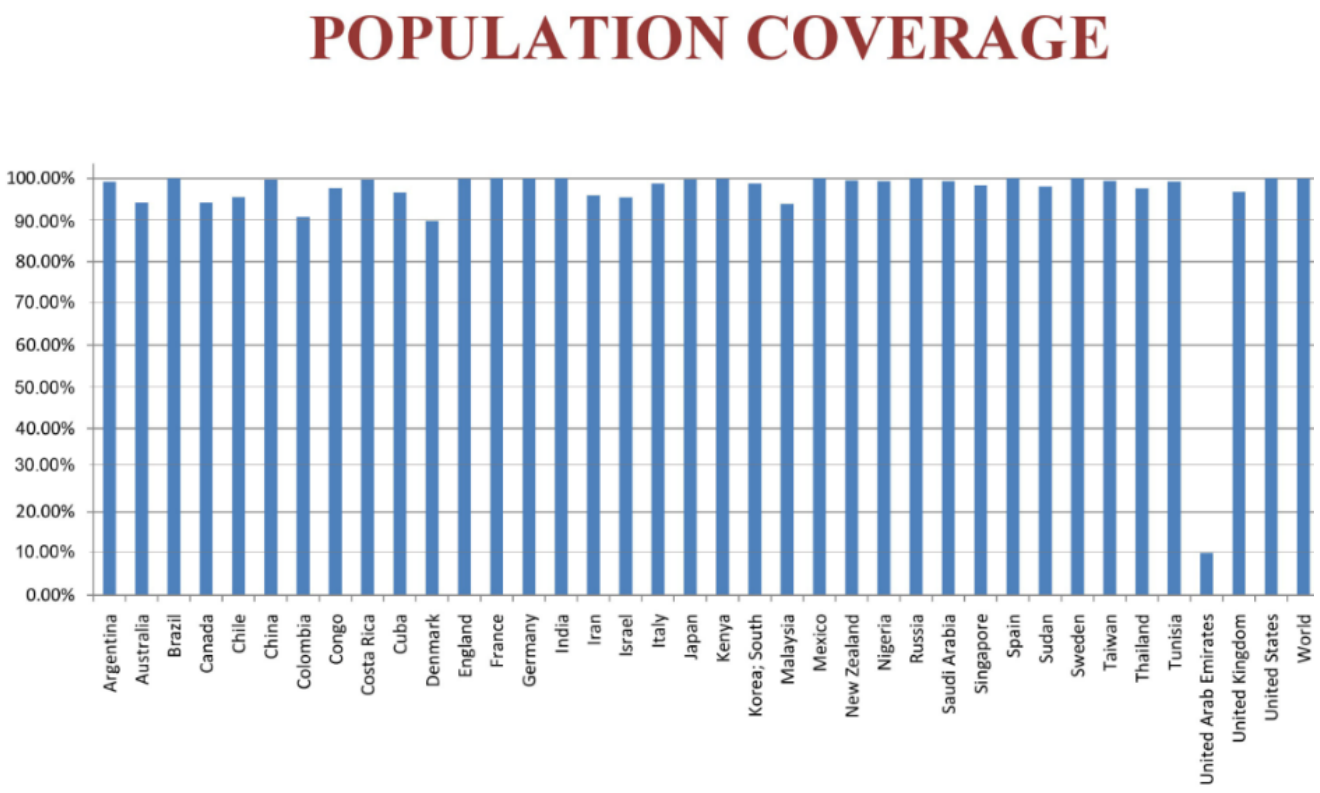


**Fig. S11. Highest and the least population coverage**

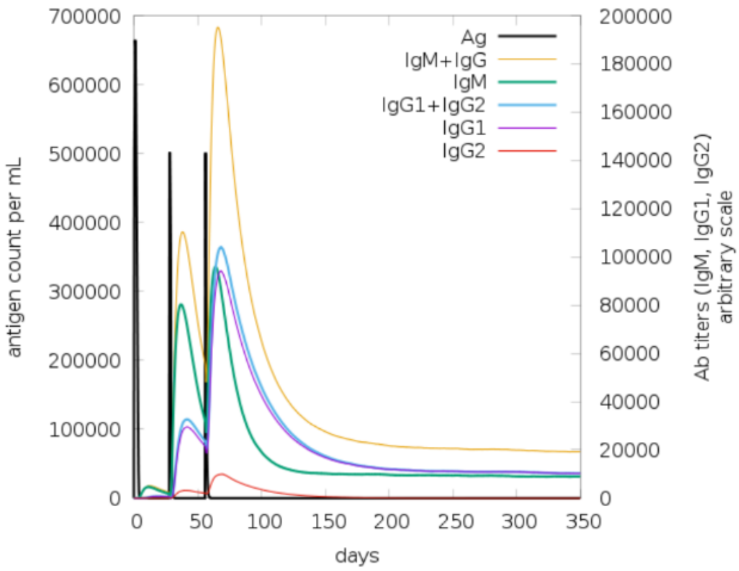


**Antigen and immunoglobulins.** Antibodies are sub-divided per isotype.

**a**

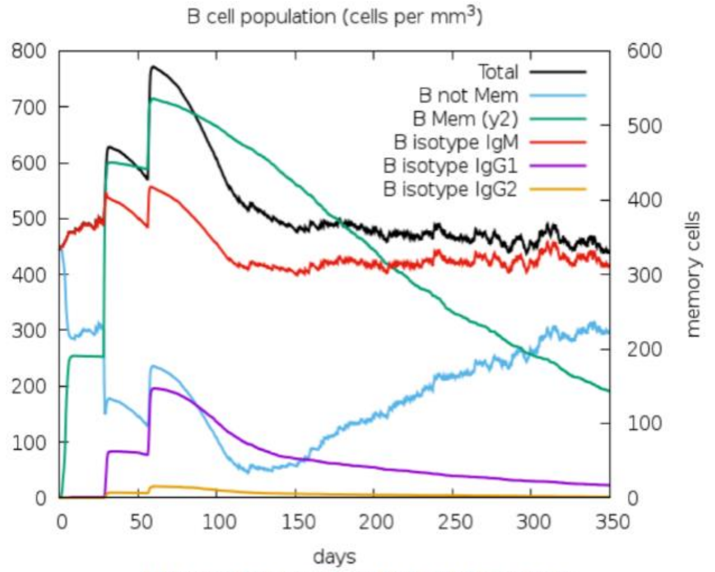


**B lymphocytes**: total count, memory cells, and sub-divided in isotypes IgM, IgG1 and IgG2.

**b**

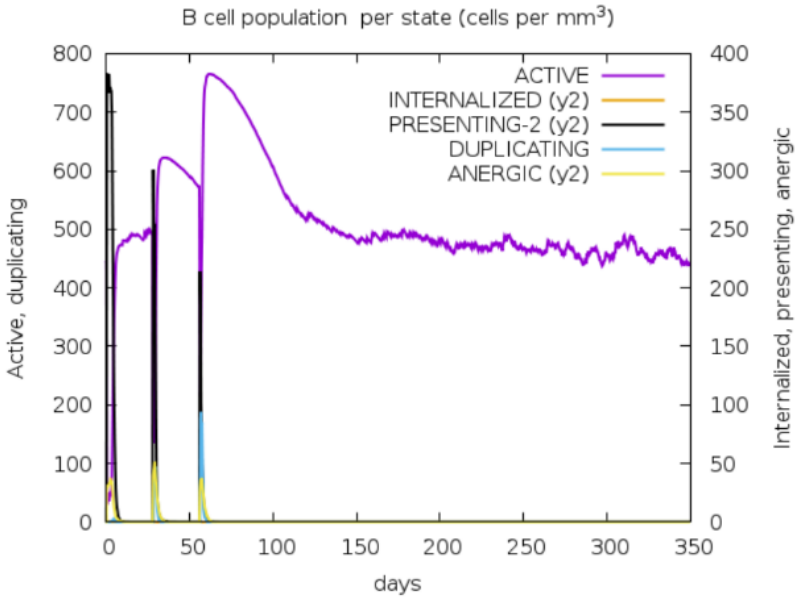


**B lymphocytes** population per entity-state (i.e., showing counts for active, presenting on class-II, internalized the Ag, duplicating and anergic.

**c**

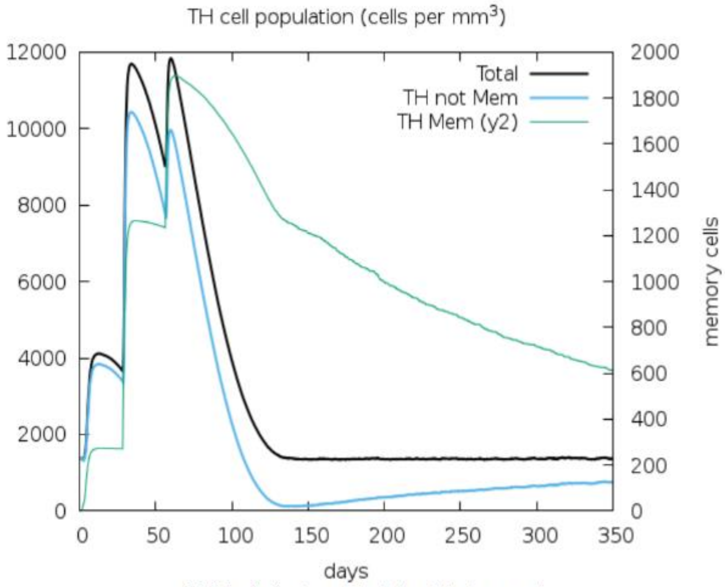


**CD4 T-helper lymphocytes** count. The plot shows total and memory counts.

**d**

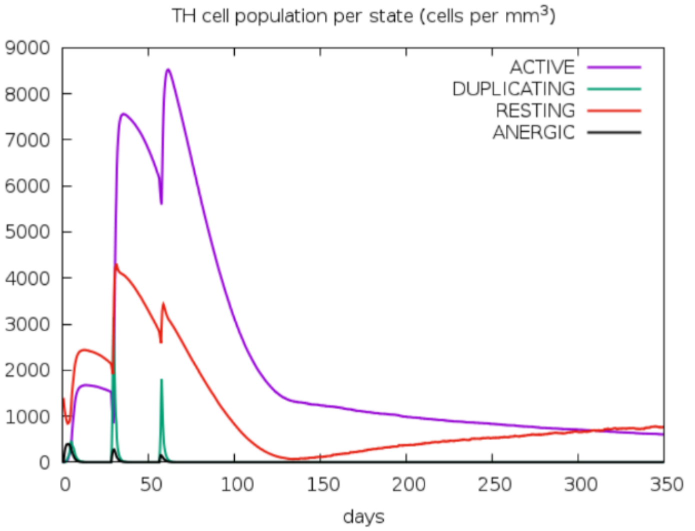


**CD4 T-helper lymphocytes** count sub-divided per entity-state (i.e., acctive, resting, anergic and duplicating).

**e**

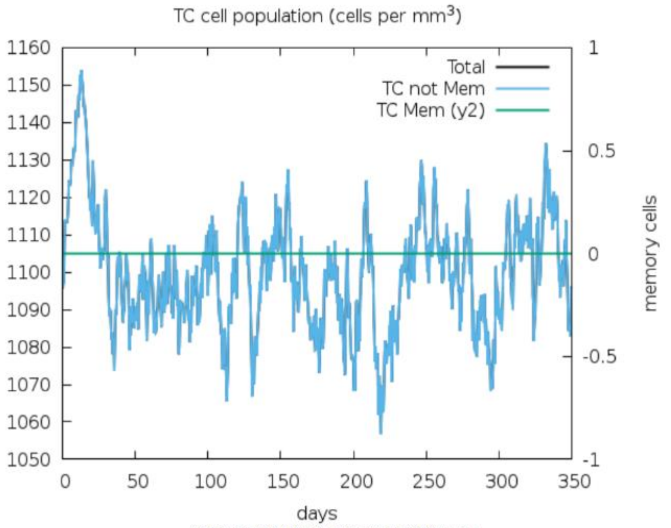


**CD8 T-cytotoxic lymphocytes** count. Total and memory shown.

**f**

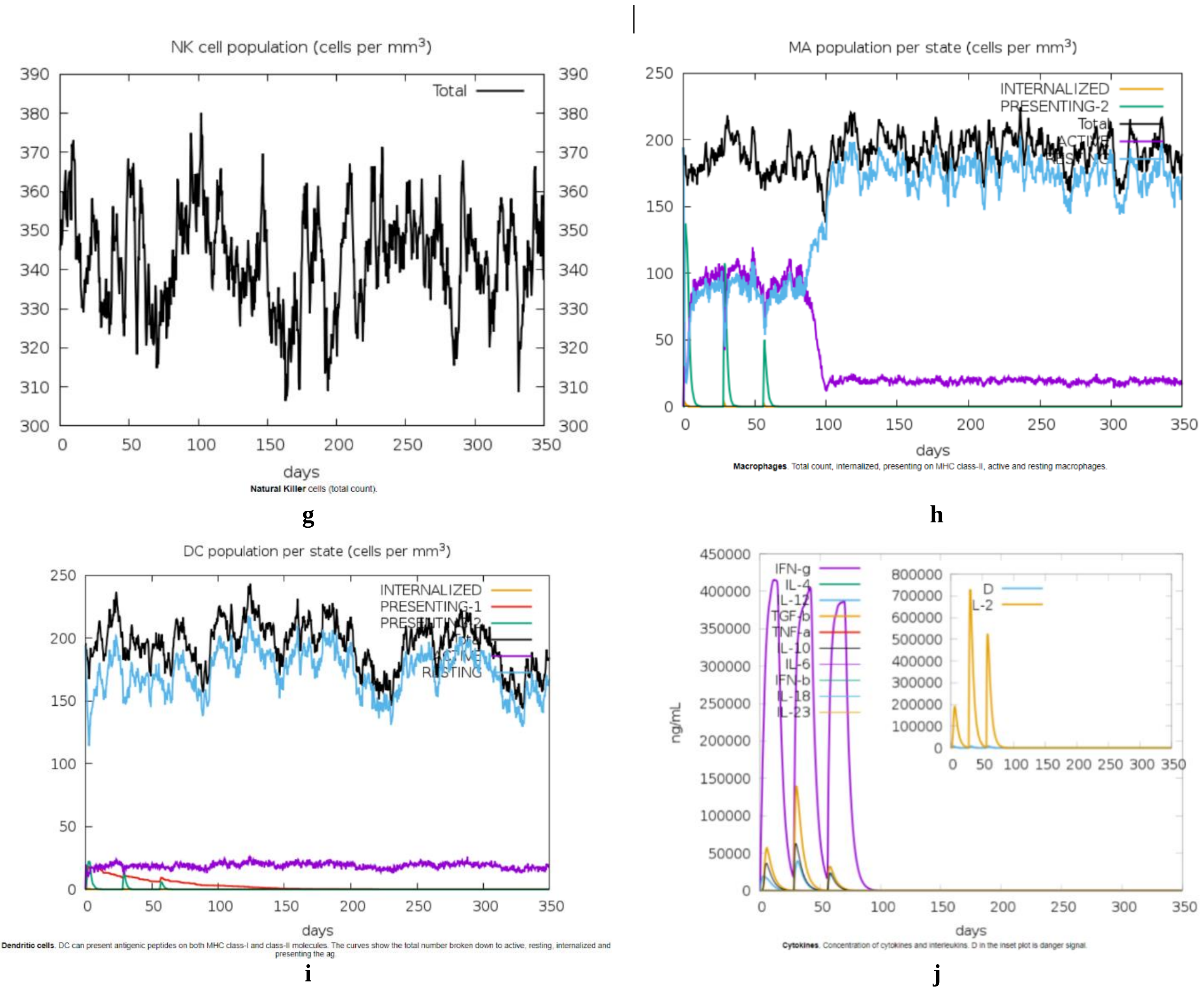


**Fig. S12. Immune simulations of vaccine construct**
