## Supplementary material for "Immunoinformatic Approach for the identification of T Cell and B Cell Epitopes in the Surface Glycoprotein and Designing a Potent Multiepitope Vaccine Construct Against SARS-CoV-2 including the new UK variant": Supplementary Tables.pdf

| Features | Value |
|---|---|
| Number of amino acids | 1273 |
| Molecular Weight | 141178.47 |
| Theoretical pI | 6.24 |
| Total number of Negatively charged residues (Asp +Glu) | 110 |
| Total number of Positively charged residues (Arg +Lys) | 103 |
| Instability Index | 33.01 |
| Aliphatic Index | 84.67 |
| Grand Average of Hydropathicity (GRAVY) | -0.079 |

**Table S1. Physiochemical properties of the spike protein of SARS-CoV-2**

| No | Epitope | Combined Score | VaxiJen Score | Remarks | No | Epitope | Combined Score | VaxiJen Score | Remarks |
|---|---|---|---|---|---|---|---|---|---|
| **1** | LTDEMIAQY | 3.6616 | 0.1043 | -- | **51** | LPPAYTNSF | 1.5189 | 0.3775 | -- |
| **2** | WTAGAAAYY | 3.1128 | 0.6306 | √ | **52** | EPVLKGVKL | 1.2802 | 1.2301 | √ |
| **3** | TSNQVAVLY | 3.0758 | 0.4387 | √ | **53** | WPWYIWLGF | 1.2753 | 1.4953 | √ |
| **4** | CVADYSVLY | 2.5759 | -0.0293 | -- | **54** | MIAQYTSAL | 1.2704 | 0.1114 | -- |
| **5** | KTSVDCTMY | 2.3795 | 1.1824 | √ | **55** | YLQPRTFLL | 1.9947 | 0.4532 | √ |
| **6** | STECSNLLL | 2.3492 | 0.4871 | √ | **56** | LLALHRSYL | 1.4391 | 0.5241 | √ |
| **7** | GAEHVNNSY | 1.9960 | 0.9347 | √ | **57** | FRKSNLKPF | 1.3950 | 0.6280 | √ |
| **8** | NIDGYFKIY | 1.9606 | -0.2462 | -- | **58** | FLHVTYVPA | 1.3772 | 1.3346 | √ |
| **9** | YSSANNCTF | 1.9531 | -0.1036 | -- | **59** | MIAQYTSAL | 1.3248 | 0.1114 | -- |
| **10** | WMESEFRVY | 1.9232 | 0.2698 | -- | **60** | GRLQSLQTY | 1.7162 | -0.0743 | -- |
| **11** | SANNCTFEY | 1.8739 | -0.0924 | -- | **61** | YRLFRKSNL | 1.6065 | 0.0522 | -- |
| **12** | VASQSIIAY | 1.7978 | 0.1366 | -- | **62** | RRARSVASQ | 1.5720 | 0.5490 | √ |
| **13** | NSFTRGVYY | 1.6915 | -0.1877 | -- | **63** | TRFQTLLAL | 1.4733 | 0.3406 | -- |
| **14** | CNDPFLGVY | 1.3355 | 0.4295 | √ | **64** | FRKSNLKPF | 1.3086 | 0.6280 | √ |
| **15** | FTNVYADSF | 1.3208 | -0.5651 | -- | **65** | YQPYRVVVL | 1.9051 | 0.5964 | √ |
| **16** | YLQPRTFLL | 1.5152 | 0.4532 | √ | **66** | TRFQTLLAL | 1.6214 | 0.3406 | -- |
| **17** | KIADYNYKL | 1.4347 | 1.6639 | √ | **67** | DEDDSEPVL | 1.3047 | 0.5104 | √ |
| **18** | SIIAYTMSL | 1.3658 | 0.5234 | √ | **68** | FEYVSQPFL | 1.2829 | 0.6324 | √ |
| **19** | VLNDILSRL | 1.3533 | -0.8524 | -- | **69** | YRLFRKSNL | 1.2575 | 0.0522 | -- |
| **20** | RLQSLQTYV | 1.2727 | -0.2167 | -- | **70** | FEYVSQPFL | 1.8668 | 0.6324 | √ |
| **21** | RLFRKSNLK | 1.7563 | -0.2829 | -- | **71** | AEIRASANL | 1.8005 | 0.7082 | √ |
| **22** | GVYFASTEK | 1.4615 | 0.7112 | √ | **72** | GEVFNATRF | 1.7314 | -0.1511 | -- |
| **23** | QIYKTPPIK | 1.4526 | -0.0833 | -- | **73** | FERDISTEI | 1.4106 | -0.7442 | -- |
| **24** | VTYVPAQEK | 1.3960 | 0.8132 | √ | **74** | AEVQIDRLI | 1.2946 | -0.5562 | -- |
| **25** | TLKSFTVEK | 1.3483 | 0.0809 | -- | **75** | DEDDSEPVL | 1.2815 | 0.5104 | √ |
| **26** | NSASFSTFK | 1.3454 | 0.1232 | -- | **76** | RSFIEDLLF | 1.9914 | -0.5782 | -- |
| **27** | KVFRSSVLH | 1.3419 | -0.6913 | -- | **77** | YSSANNCTF | 1.8837 | -0.1036 | -- |
| **28** | MTSCCSCLK | 1.3360 | 0.4270 | √ | **78** | HADQLTPTW | 1.7854 | 0.3807 | -- |

| | | | | | | | | | |
|---|---|---|---|---|---|---|---|---|---|
| **29** | GVYYHKNNK | 1.3335 | 0.8264 | √ | **79** | NSIAIPTNF | 1.6445 | 0.1744 | -- |
| **30** | KCYGVSPTK | 1.2722 | 1.4199 | √ | **80** | QSAPHGVVF | 1.6400 | 0.2234 | -- |
| **31** | NYNYLYRLF | 1.9482 | -0.1480 | -- | **81** | IAIPTNFTI | 1.5865 | 0.7052 | √ |
| **32** | PYRVVVLSF | 1.8786 | 1.0281 | √ | **82** | FAMQMAYRF | 1.5848 | 1.0278 | √ |
| **33** | VYSTGSNVF | 1.8571 | -0.3099 | -- | **83** | GTITSGWTF | 1.5724 | 0.3272 | -- |
| **34** | QYIKWPWYI | 1.8109 | 1.4177 | √ | **84** | TSNQVAVLY | 1.4503 | 0.4387 | √ |
| **35** | EYVSQPFLM | 1.7025 | 0.2605 | -- | **85** | KTSVDCTMY | 1.4355 | 1.1824 | √ |
| **36** | YFPLQSYGF | 1.6931 | 0.5107 | √ | **86** | FTNVYADSF | 1.4262 | -0.5651 | -- |
| **37** | IYQTSNFRV | 1.6294 | 0.3109 | -- | **87** | LAGTITSGW | 1.4057 | 0.3218 | -- |
| **38** | TFEYVSQPF | 1.6168 | 0.6641 | √ | **88** | SANNCTFEY | 1.3729 | -0.0924 | -- |
| **39** | VYAWNRKRI | 1.5755 | 0.5003 | √ | **89** | VFVSNGTHW | 1.2876 | 0.3438 | -- |
| **40** | AYSNNSIAI | 1.5591 | 0.8274 | √ | **90** | TLLALHRSY | 1.3654 | 0.8009 | √ |
| **41** | PFFSNVTWF | 1.4947 | 0.6638 | √ | **91** | YSSANNCTF | 1.3628 | -0.1036 | -- |
| **42** | NYLYRLFRK | 1.3274 | -0.8611 | -- | **92** | WTAGAAAYY | 1.3574 | 0.6306 | √ |
| **43** | FVFKNIDGY | 2.2795 | -0.1304 | -- | **93** | VASQSIIAY | 1.3405 | 0.1366 | -- |
| **44** | WTAGAAAYY | 2.0048 | 0.6306 | √ | **94** | QLTPTWRVY | 1.3281 | 1.2119 | √ |
| **45** | ECSNLLLQY | 1.5378 | 0.3331 | -- | **95** | GQTGKIADY | 1.3104 | 1.4019 | √ |
| **46** | ETKCTLKSF | 1.5231 | 0.8720 | √ | **96** | GVYYPDKVF | 1.2993 | 0.0652 | -- |
| **47** | CVADYSVLY | 1.5087 | -0.0293 | -- | **97** | FQPTNGVGY | 1.2969 | 0.3114 | -- |
| **48** | NGVEGFNCY | 1.4016 | 0.7783 | √ | **98** | FVSNGTHWF | 1.2938 | 0.0807 | -- |
| **49** | SPRRARSVA | 1.5619 | 0.7729 | √ | **99** | LGAENSVAY | 1.2832 | 0.4173 | √ |
| **50** | IPTNFTISV | 1.5427 | 0.8820 | √ | **100** | SVLYNSASF | 1.2612 | 0.1857 | -- |

**Table S2. NetCTL predicted epitopes and their combined score and antigenicity**

| No | Epitope | Combined Score | Vaxijen Score | Immunogenicity | Remarks |
|---|---|---|---|---|---|
| 1 | WTAGAAAYY | 3.1128 | 0.6306 | 0.15259 | √ |
| 2 | TSNQVAVLY | 3.0758 | 0.4387 | -0.01327 | √ |
| 3 | KTSVDCTMY | 2.3795 | 1.1824 | -0.11115 | √ |
| 4 | STECSNLLL | 2.3492 | 0.4871 | -0.20478 | √ |
| 5 | GAEHVNNSY | 1.9960 | 0.9347 | -0.00296 | √ |
| 6 | CNDPFLGVY | 1.3355 | 0.4295 | 0.15232 | √ |
| 7 | KIADYNYKL | 1.4347 | 1.6639 | -0.10379 | √ |
| 8 | SIIAYTMSL | 1.3658 | 0.5234 | -0.12935 | √ |
| 9 | GVYFASTEK | 1.4615 | 0.7112 | 0.09023 | √ |
| 10 | VTYVPAQEK | 1.3960 | 0.8132 | 0.02711 | √ |
| 11 | MTSCCSCLK | 1.3360 | 0.4270 | -0.36816 | √ |
| 12 | GVYYHKNNK | 1.3335 | 0.8264 | -0.18566 | √ |
| 13 | KCYGVSPTK | 1.2722 | 1.4199 | -0.06931 | √ |
| 14 | PYRVVVLSF | 1.8786 | 1.0281 | 0.03138 | √ |
| 15 | QYIKWPWYI | 1.8109 | 1.4177 | 0.21624 | √ |
| 16 | YFPLQSYGF | 1.6931 | 0.5107 | -0.26661 | √ |
| 17 | TFEYVSQPF | 1.6168 | 0.6641 | -0.19099 | √ |
| 18 | VYAWNRKRI | 1.5755 | 0.5003 | 0.12625 | √ |
| 19 | AYSNNSIAI | 1.5591 | 0.8274 | -0.08706 | √ |
| 20 | PFFSNVTWF | 1.4947 | 0.6638 | 0.06627 | √ |
| 21 | ETKCTLKSF | 1.5231 | 0.8720 | -0.37555 | √ |
| 22 | NGVEGFNCY | 1.4016 | 0.7783 | 0.22039 | √ |
| 23 | SPRRARSVA | 1.5619 | 0.7729 | 0.0402 | √ |
| 24 | IPTNFTISV | 1.5427 | 0.8820 | 0.17229 | √ |
| 25 | EPVLKGVKL | 1.2802 | 1.2301 | -0.26702 | √ |
| 26 | WPWYIWLGF | 1.2753 | 1.4953 | 0.41673 | √ |
| 27 | YLQPRTFLL | 1.9947 | 0.4532 | 0.1305 | √ |
| 28 | LLALHRSYL | 1.4391 | 0.5241 | -0.06002 | √ |

| | | | | | |
|---|---|---|---|---|---|
| 29 | FRKSNLKPF | 1.3950 | 0.6280 | -0.44169 | √ |
| 30 | FLHVTYVPA | 1.3772 | 1.3346 | 0.11472 | √ |
| 31 | RRARSVASQ | 1.5720 | 0.5490 | -0.1211 | √ |
| 32 | YQPYRVVVL | 1.9051 | 0.5964 | 0.1409 | √ |
| 33 | DEDDSEPVL | 1.3047 | 0.5104 | -0.02257 | √ |
| 34 | FEYVSQPFL | 1.8668 | 0.6324 | -0.17076 | √ |
| 35 | AEIRASANL | 1.8005 | 0.7082 | 0.00689 | √ |
| 36 | IAIPTNFTI | 1.5865 | 0.7052 | 0.18523 | √ |
| 37 | FAMQMAYRF | 1.5848 | 1.0278 | -0.28061 | √ |
| 38 | TLLALHRSY | 1.3654 | 0.8009 | 0.00244 | √ |
| 39 | QLTPTWRVY | 1.3281 | 1.2119 | 0.31555 | √ |
| 40 | GQTGKIADY | 1.3104 | 1.4019 | 0.00796 | √ |
| 41 | LGAENSVAY | 1.2832 | 0.4173 | 0.00912 | √ |

**Table S3. Predicted CTL epitopes with combined score, Antigenicity and Immunogenicity**

| Epitope | Combined Score | Vaxijen Score | Immunogenicity | Conservancy | Position | Allergenicity | Toxicity |
|---|---|---|---|---|---|---|---|
| WTAGAAAYY | 3.1128 | 0.6306 | 0.15259 | 100% | 258-266 | NA | No |
| CNDPFLGVY | 1.3355 | 0.4295 | 0.15232 | 100% | 136-144 | A | No |
| GVYFASTEK | 1.4615 | 0.7112 | 0.09023 | 100% | 89-97 | NA | No |
| VTYVPAQEK | 1.3960 | 0.8132 | 0.02711 | 100% | 1065-1073 | A | No |
| PYRVVVLSF | 1.8786 | 1.0281 | 0.03138 | 100% | 507-515 | NA | No |
| QYIKWPWYI | 1.8109 | 1.4177 | 0.21624 | 100% | 1208-1216 | A | No |
| VYAWNRKRI | 1.5755 | 0.5003 | 0.12625 | 100% | 350-358 | A | No |
| PFFSNVTWF | 1.4947 | 0.6638 | 0.06627 | 100% | 57-65 | A | No |
| NGVEGFNCY | 1.4016 | 0.7783 | 0.22039 | 100% | 481-489 | A | No |
| SPRRARSVA | 1.5619 | 0.7729 | 0.0402 | 100% | 680-688 | A | No |
| IPTNFTISV | 1.5427 | 0.8820 | 0.17229 | 100% | 714-722 | NA | No |
| WPWYIWLGF | 1.2753 | 1.4953 | 0.41673 | 98.18% | 1212-1220 | A | No |
| YLQPRTFLL | 1.9947 | 0.4532 | 0.1305 | 98.18% | 269-277 | A | No |
| FLHVTYVPA | 1.3772 | 1.3346 | 0.11472 | 100% | 1062-1070 | A | No |
| YQPYRVVVL | 1.9051 | 0.5964 | 0.1409 | 100% | 505-513 | NA | No |
| AEIRASANL | 1.8005 | 0.7082 | 0.00689 | 100% | 1016-1024 | NA | No |
| IAIPTNFTI | 1.5865 | 0.7052 | 0.18523 | 100% | 712-720 | NA | No |
| TLLALHRSY | 1.3654 | 0.8009 | 0.00244 | 98.18% | 240-248 | A | No |
| QLTPTWRVY | 1.3281 | 1.2119 | 0.31555 | 100% | 628-636 | NA | No |
| GQTGKIADY | 1.3104 | 1.4019 | 0.00796 | 100% | 413-421 | NA | No |
| LGAENSVAY | 1.2832 | 0.4173 | 0.00912 | 100% | 699-707 | NA | No |

**Table S4. Predicted CTL epitopes with combined score, Antigenicity, Immunogenicity, conservancy, Allergenicity and Toxicity (Red coloured epitopes were selected for further studies)**

| No | Epitopes | MHC1 Alleles | $IC_{50}$ (<100 nM) |
|---|---|---|---|
| 1 | WTAGAAAYY | HLA-C*03:03 | 6.82 |
| | | HLA-A*29:02 | 25.28 |
| | | HLA-C*12:03 | 52.75 |
| | | HLA-A*30:02 | 58.51 |
| | | HLA-B*15:02 | 71.82 |
| | | HLA-A*26:01 | 78.45 |
| | | HLA-B*35:01 | 83.10 |
| 2 | GVYFASTEK | HLA-C*03:03 | 11.46 |
| | | HLA-A*11:01 | 27.31 |
| | | HLA-C*12:03 | 29.05 |
| | | HLA-A*03:01 | 54.05 |
| | | HLA-C*14:02 | 91.54 |
| 3 | PYRVVVLSF | HLA-C*14:02 | 9.45 |
| | | HLA-A*23:01 | 46.71 |
| | | HLA-C*12:03 | 61.97 |
| | | HLA-C*07:02 | 66.86 |
| 4 | IPTNFTISV | HLA-C*12:03 | 18.25 |
| 5 | YQPYRVVVL | HLA-C*12:03 | 37.78 |
| | | HLA-A*02:06 | 68.16 |
| | | HLA-B*39:01 | 75.93 |
| | | HLA-B*15:02 | 92.95 |
| 6 | AEIRASANL | HLA-C*03:03 | 11.98 |
| | | HLA-B*40:01 | 25.88 |
| | | HLA-B*15:02 | 55.88 |
| | | HLA-C*12:03 | 70.02 |
| | | HLA-B*40:02 | 93.51 |
| 7 | IAIPTNFTI | HLA-C*03:03 | 6.02 |
| | | HLA-C*12:03 | 11.05 |
| | | HLA-B*58:01 | 26.63 |
| 8 | QLTPTWRVY | HLA-C*03:03 | 29.49 |
| | | HLA-C*12:03 | 44.18 |
| | | HLA-C*14:02 | 54.78 |
| | | HLA-B*15:02 | 90.20 |
| 9 | GQTGKIADY | HLA-C*03:03 | 15.69 |
| | | HLA-C*12:03 | 36.08 |
| 10 | LGAENSVAY | HLA-B*35:01 | 8.58 |
| | | HLA-C*03:03 | 20.92 |
| | | HLA-C*12:03 | 45.94 |

**Table S5. Allele selection for CTL epitopes**

| Epitope | Allele |
|---|---|
| QSLLIVNNATNVVIK | HLA-DRB1*13:02 |
| | HLA-DRB1*04:04 |
| | HLA-DRB1*04:01 |
| GINITRFQTLLALHR | HLA-DRB5*01:01 |
| | HLA-DRB1*04:04 |
| | HLA-DRB1*04:01 |
| YRVVVLSFELLHAPA | HLA-DPA1*02:01/DPB1*01:01 |
| | HLA-DPA1*01:03/DPB1*02:01 |
| | HLA-DPA1*01/DPB1*04:01 |
| FGGFNFSQILPDPSK | HLA-DRB1*04:05 |
| MFVFLVLLPLVSSQC | HLA-DRB1*01:01 |
| | HLA-DPA1*03:01/DPB1*04:02 |
| VLSFELLHAPATVCG | HLA-DRB1*04:05 |
| | HLA-DRB1*04:04 |
| | HLA-DRB1*11:01 |
| | HLA-DPA1*01:03/DPB1*02:01 |
| TESIVRFPNITNLCP | HLA-DRB1*04:04 |
| ITSGWTFGAGAALQI | HLA-DRB1*09:01 |
| | HLA-DQA1*05:01/DQB1*03:01 |
| QDLFLPFFSNVTWFH | HLA-DRB1*15:01 |
| | HLA-DPA1*01/DPB1*04:01 |
| AIPTNFTISVTTEIL | HLA-DRB1*07:01 |
| CSNLLLQYGSFCTQL | HLA-DRB1*15:01 |
| YIWLGFIAGLIAIVM | HLA-DQA1*05:01/DQB1*03:01 |
| | HLA-DRB1*12:01 |
| RFASVYAWNRKRISN | HLA-DRB1*11:01 |
| | HLA-DRB5*01:01 |
| GLTVLPPLLTDEMIA | HLA-DRB1*04:04 |

**Table S6. Allele selection for HTL epitopes**

Predicted Discontinuous Epitope(s):

| No. | Residues | Number of residues | Score | 3D structure |
|---|---|---|---|---|
| 1 | A:D1139, A:P1140, A:L1141, A:Q1142, A:P1143, A:E1144, A:L1145, A:D1146 | 8 | 0.97 | View |
| 2 | A:N343, A:A344, A:T345, A:R346, A:F347, A:A348, A:S349, A:V350, A:Y351, A:A352, A:W353, A:N354, A:S399, A:V401, A:I402, A:R403, A:G404, A:T415, A:G416, A:K417, A:I418, A:A419, A:D420, A:Y421, A:N422, A:Y423, A:K424, A:S438, A:N439, A:N440, A:L441, A:D442, A:S443, A:K444, A:V445, A:G446, A:G447, A:N448, A:Y449, A:N450, A:Y451, A:L452, A:Y453, A:R454, A:L455, A:F456, A:R457, A:K458, A:S459, A:N460, A:L461, A:K462, A:P463, A:F464, A:E465, A:R466, A:D467, A:I468, A:S469, A:T470, A:E471, A:I472, A:Y473, A:Q474, A:A475, A:G476, A:S477, A:T478, A:P479, A:C480, A:N481, A:G482, A:V483, A:E484, A:G485, A:F486, A:N487, A:C488, A:Y489, A:F490, A:P491, A:L492, A:Q493, A:S494, A:Y495, A:G496, A:F497, A:Q498, A:P499, A:T500, A:N501, A:G502, A:V503, A:G504, A:Y505, A:Q506, A:P507, A:R509 | 98 | 0.894 | View |
| 3 | A:Y707, A:S708, A:N709, A:N710, A:S711, A:F1075, A:T1076, A:T1077, A:A1078, A:P1079, A:A1080, A:I1081, A:C1082, A:H1083, A:D1084, A:G1085, A:K1086, A:A1087, A:H1088, A:F1089, A:P1090, A:R1091, A:E1092, A:G1093, A:V1094, A:F1095, A:V1096, A:S1097, A:N1098, A:G1099, A:T1100, A:H1101, A:W1102, A:F1103, A:V1104, A:R1107, A:Y1110, A:E1111, A:P1112, A:Q1113, A:I1114, A:I1115, A:T1116, A:T1117, A:D1118, A:N1119, A:T1120, A:F1121, A:V1122, A:S1123, A:G1124, A:N1125, A:C1126, A:D1127, A:V1128, A:V1129, A:I1130, A:G1131, A:I1132, A:V1133, A:N1134, A:N1135, A:T1136, A:V1137, A:Y1138 | 65 | 0.887 | View |
| 4 | A:H66, A:A67, A:I68, A:H69, A:V70, A:S71, A:G72, A:T73, A:N74, A:G75, A:T76, A:K77, A:R78, A:E96, A:K97, A:S98, A:N99, A:I100, A:D138, A:P139, A:F140, A:L141, A:G142, A:V143, A:Y144, A:Y145, A:H146, A:K147, A:N148, A:N149, A:K150, A:S151, A:W152, A:M153, A:E154, A:S155, A:E156, A:F157, A:R158, A:V159, A:Y160, A:A243, A:L244, A:H245, A:R246, A:S247, A:Y248, A:L249, A:T250, A:P251, A:G252, A:D253, A:S254, A:S255, A:S256, A:G257, A:W258, A:T259, A:A260, A:G261, A:A262, A:A263 | 62 | 0.872 | View |
| 5 | A:N122, A:A123, A:T124 | 3 | 0.815 | View |
| 6 | A:D178, A:L179, A:E180, A:G181, A:K182, A:K187 | 6 | 0.806 | View |

**Table S7. Conformational /discontinuous B cell epitope prediction**

| Cluster | Members | Representative | Weighted Score |
|---|---|---|---|
| **0** | 47 | Center | -1050.4 |
| | | Lowest Energy | -1274.5 |
| **1** | 34 | Center | -1052.0 |
| | | Lowest Energy | -1157.0 |
| **2** | 25 | Center | -1221.0 |
| | | Lowest Energy | -1221.0 |
| **3** | 22 | Center | -1070.0 |
| | | Lowest Energy | -1145.1 |
| **4** | 21 | Center | -968.5 |
| | | Lowest Energy | -1169.4 |
| **5** | 21 | Center | -1154.2 |
| | | Lowest Energy | -1154.2 |
| **6** | 21 | Center | -1076.5 |
| | | Lowest Energy | -1134.7 |
| **7** | 19 | Center | -1106.3 |
| | | Lowest Energy | -1106.3 |
| **8** | 18 | Center | -1079.3 |
| | | Lowest Energy | -1079.3 |
| **9** | 18 | Center | -948.2 |
| | | Lowest Energy | -998.0 |
| **10** | 17 | Center | -1014.7 |
| | | Lowest Energy | -1115.7 |
| **11** | 17 | Center | -1069.0 |
| | | Lowest Energy | -1069.0 |
| **12** | 17 | Center | -949.9 |
| | | Lowest Energy | -1185.2 |
| **13** | 16 | Center | -1158.3 |
| | | Lowest Energy | -1158.3 |
| **14** | 16 | Center | -1044.5 |
| | | Lowest Energy | -1181.4 |
| **15** | 14 | Center | -942.5 |
| | | Lowest Energy | -1022.7 |
| **16** | 14 | Center | -941.9 |
| | | Lowest Energy | -1115.6 |

| Cluster | Members | Representative | Weighted Score |
|---|---|---|---|
| **17** | 14 | Center | -957.7 |
| | | Lowest Energy | -988.3 |
| **18** | 13 | Center | -1041.8 |
| | | Lowest Energy | -1041.8 |
| **19** | 12 | Center | -1202.5 |
| | | Lowest Energy | -1202.5 |
| **20** | 12 | Center | -1060.3 |
| | | Lowest Energy | -1060.3 |
| **21** | 12 | Center | -954.6 |
| | | Lowest Energy | -997.2 |
| **22** | 12 | Center | -949.2 |
| | | Lowest Energy | -1139.1 |
| **23** | 11 | Center | -980.1 |
| | | Lowest Energy | -1097.8 |
| **24** | 11 | Center | -978.8 |
| | | Lowest Energy | -1081.9 |
| **25** | 11 | Center | -942.5 |
| | | Lowest Energy | -1094.0 |
| **26** | 11 | Center | -1143.4 |
| | | Lowest Energy | -1161.5 |
| **27** | 11 | Center | -1107.3 |
| | | Lowest Energy | -1107.3 |
| **28** | 11 | Center | -1106.0 |
| | | Lowest Energy | -1106.0 |
| **29** | 11 | Center | -1075.4 |
| | | Lowest Energy | -1075.4 |

**Table S8. TLR3 Cluspro scores**

| Cluster | Members | Representative | Weighted Score |
|---|---|---|---|
| **0** | 37 | Center | -1166.9 |
| | | Lowest Energy | -1329.1 |
| **1** | 31 | Center | -1189.3 |
| | | Lowest Energy | -1189.3 |
| **2** | 29 | Center | -1103.8 |
| | | Lowest Energy | -1156.2 |
| **3** | 26 | Center | -1138.5 |
| | | Lowest Energy | -1138.5 |
| **4** | 24 | Center | -1067.6 |
| | | Lowest Energy | -1164.3 |
| **5** | 23 | Center | -1089.7 |
| | | Lowest Energy | -1194.0 |
| **6** | 23 | Center | -1136.6 |
| | | Lowest Energy | -1205.6 |
| **7** | 19 | Center | -1002.0 |
| | | Lowest Energy | -1143.3 |
| **8** | 18 | Center | -1139.1 |
| | | Lowest Energy | -1191.6 |
| **9** | 18 | Center | -1108.1 |
| | | Lowest Energy | -1137.9 |
| **10** | 18 | Center | -1056.3 |
| | | Lowest Energy | -1076.1 |
| **11** | 17 | Center | -1058.4 |
| | | Lowest Energy | -1366.7 |
| **12** | 15 | Center | -1111.2 |
| | | Lowest Energy | -1233.2 |
| **13** | 14 | Center | -1023.1 |
| | | Lowest Energy | -1132.8 |
| **14** | 13 | Center | -980.9 |
| | | Lowest Energy | -1191.6 |
| **15** | 13 | Center | -1019.1 |
| | | Lowest Energy | -1168.4 |
| **16** | 13 | Center | -1028.1 |
| | | Lowest Energy | -1101.1 |

| Cluster | Members | Representative | Weighted Score |
|---|---|---|---|
| **17** | 13 | Center | -1040.7 |
| | | Lowest Energy | -1129.2 |
| **18** | 13 | Center | -1027.9 |
| | | Lowest Energy | -1113.6 |
| **19** | 12 | Center | -1110.0 |
| | | Lowest Energy | -1139.4 |
| **20** | 12 | Center | -1070.5 |
| | | Lowest Energy | -1097.1 |
| **21** | 12 | Center | -1016.9 |
| | | Lowest Energy | -1118.2 |
| **22** | 11 | Center | -1060.2 |
| | | Lowest Energy | -1155.9 |
| **23** | 11 | Center | -1223.8 |
| | | Lowest Energy | -1223.8 |
| **24** | 11 | Center | -1072.7 |
| | | Lowest Energy | -1072.7 |
| **25** | 11 | Center | -1042.8 |
| | | Lowest Energy | -1042.8 |
| **26** | 10 | Center | -1008.2 |
| | | Lowest Energy | -1088.2 |
| **27** | 10 | Center | -984.6 |
| | | Lowest Energy | -1090.9 |
| **28** | 10 | Center | -1068.8 |
| | | Lowest Energy | -1068.8 |
| **29** | 10 | Center | -985.3 |
| | | Lowest Energy | -1078.6 |

**Table S9. TLR4 Cluspro scores**

| Population/area | Class Combined | | |
|---|---|---|---|
| | Coverage[a] | Average_hit[b] | Pc90[c] |
| Algeria | 87.06% | 2.78 | 0.77 |
| American Samoa | 87.8% | 2.23 | 0.82 |
| Argentina | 99.16% | 5.38 | 2.99 |
| Australia | 94.16% | 5.14 | 1.44 |
| Austria | 99.7% | 5.59 | 3.4 |
| Belarus | 43.81% | 0.94 | 0.36 |
| Belgium | 98.99% | 4.83 | 2.66 |
| Bolivia | 70.4% | 2.01 | 0.34 |
| Borneo | 70.34% | 1.69 | 0.34 |
| Brazil | 100.0% | 8.8 | 5.08 |
| Bulgaria | 82.76% | 2.95 | 0.58 |
| Cameroon | 99.98% | 7.02 | 4.22 |
| Canada | 94.17% | 3.44 | 2.1 |
| Cape Verde | 93.61% | 2.74 | 1.21 |
| Central African Republic | 99.01% | 2.27 | 1.41 |
| Chile | 95.5% | 4.81 | 1.65 |
| China | 99.66% | 7.78 | 3.92 |
| Colombia | 90.78% | 3.23 | 1.06 |
| Congo | 97.63% | 3.64 | 1.76 |

| | | | |
|---|---|---|---|
| Cook Islands | 100.0% | 7.86 | 5.33 |
| Costa Rica | 99.69% | 4.25 | 3.07 |
| Croatia | 94.52% | 3.6 | 1.58 |
| Cuba | 96.55% | 3.35 | 1.52 |
| Czech Republic | 99.91% | 7.21 | 4.26 |
| Denmark | 89.77% | 2.79 | 0.98 |
| Ecuador | 99.85% | 5.18 | 2.88 |
| England | 99.85% | 7.88 | 4.16 |
| Ethiopia | 88.18% | 2.47 | 0.85 |
| Fiji | 97.22% | 4.29 | 2.34 |
| Finland | 99.62% | 7.22 | 3.24 |
| France | 100.0% | 10.88 | 6.58 |
| Gabon | 99.9% | 5.4 | 3.35 |
| Gambia | 99.95% | 5.42 | 3.41 |
| Georgia | 99.6% | 8.42 | 4.08 |
| Germany | 99.91% | 8.81 | 4.67 |
| Greece | 99.26% | 5.08 | 2.85 |
| Guatemala | 41.27% | 1.04 | 0.17 |
| Guinea-Bissau | 89.75% | 2.41 | 0.98 |
| Hong Kong | 83.36% | 1.93 | 0.6 |
| India | 99.96% | 8.43 | 4.86 |
| Indonesia | 95.81% | 4.3 | 1.78 |

| | | | |
|---|---|---|---|
| Iran | 95.91% | 5.35 | 1.85 |
| Ireland Northern | 99.92% | 7.69 | 4.3 |
| Ireland South | 99.83% | 7.38 | 3.84 |
| Israel | 95.43% | 4.67 | 1.71 |
| Italy | 98.72% | 7.61 | 2.99 |
| Ivory Coast | 15.33% | 0.27 | 0.12 |
| Jamaica | 64.73% | 1.64 | 0.28 |
| Japan | 99.74% | 8.11 | 4.12 |
| Jordan | 84.53% | 3.45 | 0.65 |
| Kenya | 99.86% | 6.0 | 3.55 |
| Kiribati | 68.93% | 1.57 | 0.32 |
| Korea; South | 98.71% | 6.25 | 2.49 |
| Lebanon | 99.28% | 7.06 | 3.03 |
| Liberia | 99.5% | 2.75 | 2.02 |
| Macedonia | 92.58% | 3.6 | 1.49 |
| Malaysia | 93.91% | 3.76 | 1.44 |
| Mali | 74.19% | 1.89 | 0.39 |
| Martinique | 83.08% | 2.15 | 0.59 |
| Mexico | 100.0% | 8.56 | 5.33 |
| Mongolia | 99.4% | 5.53 | 3.16 |
| Morocco | 94.57% | 3.95 | 1.4 |
| Netherlands | 98.97% | 4.95 | 2.64 |

| | | | |
|---|---|---|---|
| New Caledonia | 98.87% | 5.95 | 3.0 |
| New Zealand | 99.51% | 5.81 | 3.53 |
| Nigeria | 99.35% | 2.97 | 2.01 |
| Niue | 98.97% | 5.43 | 3.08 |
| Norway | 99.74% | 5.71 | 3.43 |
| Oman | 67.5% | 1.06 | 0.31 |
| Pakistan | 80.61% | 2.89 | 0.52 |
| Papua New Guinea | 99.99% | 10.38 | 6.28 |
| Paraguay | 44.79% | 1.03 | 0.36 |
| Peru | 83.35% | 2.9 | 0.6 |
| Philippines | 92.97% | 3.88 | 1.16 |
| Poland | 98.12% | 6.74 | 2.34 |
| Portugal | 95.28% | 5.02 | 1.55 |
| Romania | 65.19% | 1.08 | 0.29 |
| Russia | 100.0% | 10.73 | 6.67 |
| Rwanda | 74.66% | 1.65 | 0.39 |
| Samoa | 98.98% | 5.68 | 3.14 |
| Saudi Arabia | 99.33% | 5.41 | 2.85 |
| Scotland | 93.62% | 4.91 | 1.33 |
| Senegal | 84.9% | 2.11 | 0.66 |
| Serbia | 25.7% | 0.44 | 0.13 |
| Singapore | 98.3% | 5.31 | 2.45 |

| | | | |
|---|---|---|---|
| Slovakia | 98.0% | 3.18 | 2.02 |
| Slovenia | 100.0% | 7.57 | 5.29 |
| South Africa | 76.99% | 2.08 | 0.43 |
| Spain | 100.0% | 9.38 | 4.95 |
| Sri Lanka | 23.96% | 0.24 | 0.13 |
| Sudan | 98.01% | 6.35 | 2.42 |
| Sweden | 100.0% | 8.8 | 6.2 |
| Switzerland | 56.31% | 2.54 | 0.23 |
| Taiwan | 99.42% | 6.75 | 3.27 |
| Thailand | 97.61% | 5.21 | 2.09 |
| Tokelau | 99.86% | 5.75 | 4.16 |
| Tonga | 98.03% | 4.91 | 2.57 |
| Tunisia | 99.23% | 6.56 | 3.07 |
| Turkey | 94.22% | 5.03 | 1.81 |
| Uganda | 99.93% | 5.93 | 3.31 |
| Ukraine | 50.64% | 1.1 | 0.41 |
| United Arab Emirates | 10.13% | 0.14 | 0.11 |
| United Kingdom | 96.74% | 4.65 | 2.12 |
| United States | 100.0% | 10.23 | 6.56 |
| Venezuela | 99.49% | 3.61 | 2.12 |
| Vietnam | 96.11% | 4.89 | 1.68 |
| Wales | 1.0% | 0.01 | 0.1 |

| | | | |
|---|---|---|---|
| World | 99.9% | 8.91 | 4.69 |
| Zambia | 76.21% | 1.36 | 0.42 |
| Zimbabwe | 99.46% | 4.85 | 2.58 |

[a] projected population coverage
[b] average number of epitope hits / HLA combinations recognized by the population
[c] minimum number of epitope hits / HLA combinations recognized by 90% of the population

**Table S10. Population coverage of MEVC**
